# Activity-driven emergence of genealogical enclaves in growing bacterial colonies

**DOI:** 10.1101/2023.09.07.556749

**Authors:** Garima Rani, Anupam Sengupta

## Abstract

Bacterial dispersal, the movement of cells spanning diverse physical scales and environments, has been long investigated owing to its far-reaching ramifications in the ecology and evolution of bacterial species and their consortia. A major proportion of bacterial species are surface associated, yet if and how they disperse, specifically during the early stages of biofilm formation, remains to be understood. While physical vectors like fluid flow drive dispersal across large scales, surface-associated cells may benefit from the active biomechanical forces to navigate locally within a colony. Here, by analyzing sessile bacterial colonies, we study how cells disperse over generations due to the growth-induced forces under different conditions. A custom-built labelfree algorithm, developed to track the progeny cells as they grow and divide, reveals the emergence of distinct self-similar genealogical enclaves which intermix over time. Biological activity, indicated by the division times, is a key determinant of the intermixing dynamics; while topological defects appearing at the interface of the enclaves mediate the morphology of finger-like interfacial domains. By quantifying the Shannon entropy, we show that dividing bacterial cells have spatial affinity to close relatives, at the cost of the entropically favourable option of intermixing, wherein faster growing colonies show higher drop in the Shannon entropy over time. A coarse-grained lattice modelling of such colonies, combined with insights from the thermodynamics of phase separation, suggest that the emergence of genealogical enclaves results from an interplay of growth-induced cell dispersal within the colony (which promotes intermixing) and stochasticity of cell division, alongwith the cell-cell interactions at a given growth condition. Our study uncovers the evolution of so-far hidden emergent self-organising features within growing bacterial colonies, which while displaying a high degree of self-similarity on a range of phenotypic traits, point at competing roles of growth-induced forces and entropic landscapes which ultimately shape the genealogical distance of cells to their kith and kin within growing colonies.

**One Sentence Summary:** Label-free tracking of cells reveals spatio-temporal evolution of genealogical enclaves in growing bacterial colonies resulting from activity-controlled emergent demixing of cell lineages.

## Introduction

Microbial colonies of one or several organisms are ubiquitous in nature virtually since the appearance of life on earth. Due to their relevance from ecological and public health perspectives, such agglomerations have been widely studied [1, 2, 3]. In particular, several seminal insights have been obtained on a range of topics, including cell communication strategies in microbial colonies [4, 5], collective behaviour of cells [6, 7], response to stress [8, 9]. Nevertheless, biophysical principles underpinning emergent organisation of cells in growing colonies and the role of of biological activity in driving them, remain poorly understood.

Cells in confluent bacterial colonies show several remarkable features of self organisation [10, 11, 12, 13] and the importance of structural order and topological attributes of cell organisation in optimising growth of such colonies has been highlighted [14, 15]. In particular, singularities in local ordering of cells in colonies, referred to as topological defects, have been implicated in driving the formation of layers in biofilms for both motile and sessile bacteria [16, 17], which marks a significant step in biofilm growth and maturation. Topological defects have also been shown to regulate several other fundamental features of biofilm growth and propagation like navigation [18], sporulation [19] and nutrient uptake [20]. Such defects are common in the periphery of bacterial colonies driving advancing fronts but they are also regularly observed in the interior of the colonies, seemingly at random [21]. Further, as with any structured living community, a key determinant of its resilience are interactions fostered between its constituents [22, 23]. Arguably, the most meaningful interactions are typically between close relatives on one hand and spatial neighbours on the other hand [24, 25]. Therefore, a deep understanding of the growth of microbial communities and devising of effective strategies to limit biofilm growth on demand is predicated upon the depth of our understanding of the way cells organise themselves spatially and in relation to their lineage kins in these communities. For such studies, it is essential to be able to track cells as they grow and divide over generations to form progeny chains. The primary method for tracking of cells in microbial consortia has been labelling of appropriate cells, typically by usage fluorescent proteins (FPs) [26]. Labelling of cells using FPs has been used widely and has resulted in revelation of several significant features of spatial organisation of cells in microbial colonies, over and above their visible topological features [13, 27, 28, 29]. Nevertheless, such labelling methods involve fairly sophisticated experimental manipulation, underscoring the need for developing automated, label-free cell tracking techniques.

## Emergence of genealogical enclaves in growing bacterial colonies

Here, to discern the emergent spatial structure of the genealogical organisation in growing bacterial colonies, we tracked and studied intermixing dynamics of descendants of individual cells in colonies of surface associated bacterial species, specifically *Escherichia coli*. For this, we developed a novel label-free tracking algorithm to spatially trace the progeny chains emanating from the two daughter cells arising from the first division event of the colony, that of the founder cell (Fig. 1A and Supplementary Fig. 1 and 2). This allows us to bypass relatively tedious experimental methods of tracking. Indeed, while such tracking has typically been carried out by experimental labelling based methods and usually involving different species or a combination of wild and mutant type cells, here we carry this out in an entirely label-free manner by leveraging a tracking algorithm that effectively logs the positions of cells as they grow and recognises division events so as to spatially map out the progeny chain starting from an initial cell (Fig. 1B and Supplementary Sec. S.3). For the current study, we limit our discussions to the monolayer configuration of the bacterial colony, *i.e.,* before the colony undergoes the mono-to-multilayer transition [30]. The cells of the two progeny chains undergo spatial intermixing as the colony grows, resulting in a dynamic partitioning of the colony into two domains. Since the colony grows freely on the substrate, it is natural to expect a high order of intermixing amongst the cells belonging to the two domains, which also will be entropically more favourable. However, we observed that as the colonies grow, cells from the two domains arranged themselves into enclaves, maintaining a spatial affinity for their close kin (Fig. 1A and Supplementary Fig. 2), having cell arrangement patterns very similar to the case of merging of colonies, whereby two smaller colonies merge to form a larger colony and in which case, it is more plausible and indeed observed, that the cells from the two initial colonies form enclaves within the larger colony upon merging, as its grows further (Supplementary Fig. 3).

**Figure 1:**
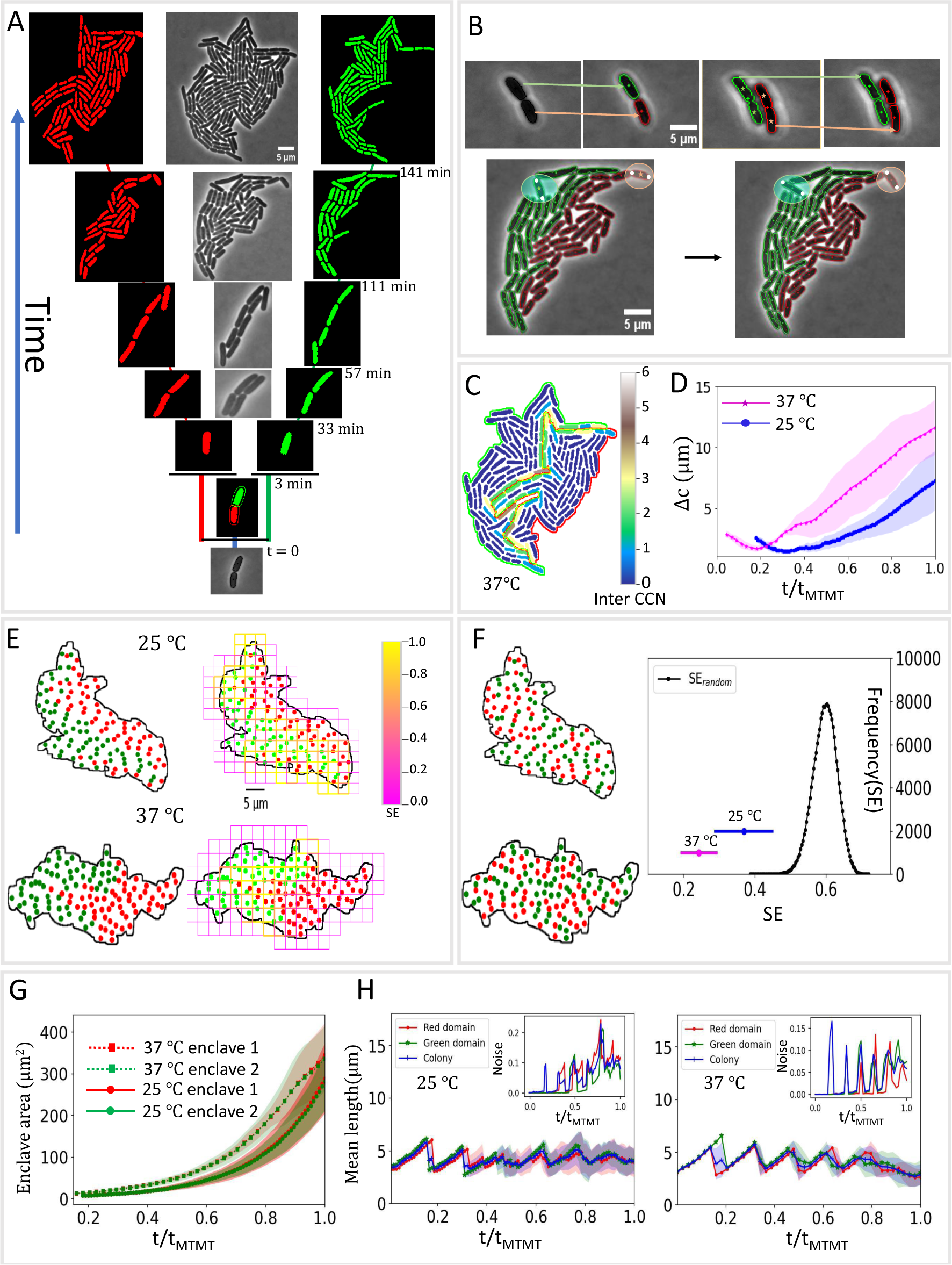
Formation of self similar enclaves in bacterial colonies. **A.** Label-free tracking of a bacterial colony reveals partitioning of the colony into lineage enclaves starting from the initial two cells following the division of the founder cell. **B.** Label-free tracking algorithm relies on frame to frame mapping of the centroid of cells to track them as they grow, with additional parameters based on mapping the cell poles and approximate prediction of daughter cell centroids to identify recognise left out cells and division events. **C.** Cell contact number (CCN) analysis to calculate neighbours of cells belonging to different progeny chain reveals that most of the cells only have spatial neighbours belonging to their own progeny chain. **D.** The relative distance (Δ*c*) between the centroids of the two enclaves is mapped as a function of time. The error (s.d.) is shown by the shaded region in the plot. **E.** Box counting method employed to calculate Shannon entropy of cell arrangement patterns in colonies. **F.** The ordered nature of enclave arrangement pattern *vis-a-vis* random arrangement patterns where progeny chain membership is assigned randomly to cells in the colony fixing colony geometry and proportion of cells in each domain (Black curve maps the values of Shannon entropy (SE) for random arrangement patterns of cells in colonies, while the Shannon entropy values for enclave arrangements for cells growing at 25 *^°^*C and 37 *^°^*C are marked in blue and magenta, respectively). **G.** The area of the two enclaves for colonies growing as a function of normalised time. **H.** Mean cell length of cells in each of the two enclaves (colored red and green) as well as the whole colony (colored blue) is mapped as a function of normalised time for cells grown at 25 *^°^*C and 37 *^°^*C (Insetphenotypic noise for cell length (quantified by the normalized variance) is plotted for the two enclaves (red and green) and the colony (blue) as a function of normalised time.

Formation of genealogical enclaves suggests that for a given cell in the colony, the majority of its neighbours will be from its progeny chain. To check this, we performed contact number analysis of cells in the colony (Supplementary Sec. S.4) and observed that indeed, the large majority of cells in the colony have very few contacts with cells not in their progeny chain and in fact, such contacts decreased as the colony matures (Fig. 1C, Fig. 2B and Supplementary Fig. 4).

**Figure 2:**
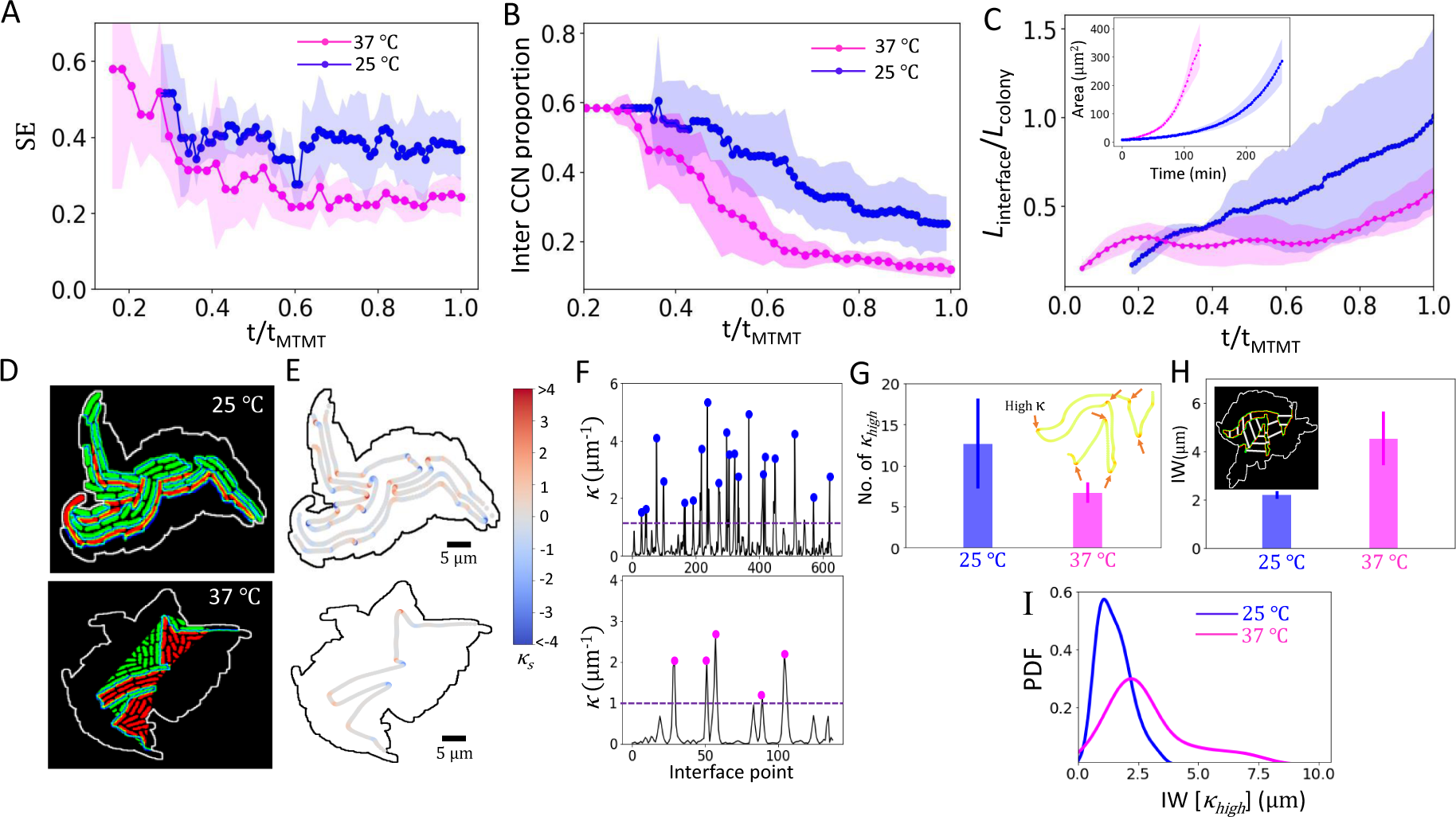
Activity mediates emergent partitioning of bacterial cells into genealogical enclaves. **A.** Shannon entropy of cell arrangement patterns emerging from enclave partitioning of colony as a function of normalised time for cells growing at 25 *^°^*C (blue) and 37 *^°^*C (magenta). **B.** Proportion of inter-enclave contacts of cells in the colony is plotted as a function of normalised time for cells growing at 25 *^°^*C (blue) and 37 *^°^*C (magenta). **C.** The length of the interface between the two enclaves (*L_interface_*) normalised by the perimeter of the entire colony (*L_colony_*) is mapped as a function of normalised time for colony growing at 25 *^°^*C (blue) and 37 C (magenta), (Inset)-The area of the two enclaves is mapped for colonies growing at 25 *^°^*C (blue) and 37 *^°^*C (magenta). **D.** The enclave invasion fronts in the colonies are highlighted in two representative cases each for cells growing at 25 *^°^*C (left) and 37 *^°^*C (right). **E.** The signed curvature (*κ_s_*) of interfacial curves in two representative cases is shown here and **F.** (modulus of) curvature (*κ*) values are mapped parametrised by the interfacial curve, with peak values of curvature above a threshold value (dotted line) highlighted by blue dots (for cells growing at 25 *^°^*C) and magenta dots (cells growing at 37 *^°^*C). **G.** Mean ± s.d. of the frequency of high curvature (*κ_high_*) points in the interfacial curves is shown as a function of temperature. **H.** Mean invasion width (IW) is plotted in the two cases of colonies growing at 25 *^°^*C (blue) and 37 *^°^*C (magenta). **I.** Probability distribution (PDF) of invasion width in the vicinity of high curvature regions (IW[*κ_high_*]) of the interfacial curve.

To quantify the spatial distribution of cells belonging to the two progeny chains relative to each other, we calculated the centroid of the two enclaves (Supplementary Sec. S.5) and the evolution of their distance from each other. While in case cells from the two progeny chains were well intermixed, the two centroid would have been close to each other, we observed the opposite, that the two centroid move farther and farther away with time (Fig. 1D). Thus, we can conclusively infer that cells in progeny chains emanating from the first two daughter cells preferentially arrange themselves into enclaves within the colony. We also observe similar formation of progeny enclaves when we varied substrate properties by varying agarose concentration (Supplementary Fig. 5) as well for *Vibrio cholerae* colonies, in which case cells have a much lower aspect ratio (Supplementary Fig. 6), thus highlighting the robustness of this phenomena under a range of conditions and for various species of bacteria.

Such dynamic partitioning of the colony into two domains gives rise to characteristic cell arrangement patterns. To quantify the level of intermixing of such patterns *vis-a-vis* randomly intermixed distributions, we calculated the Shannon entropy [31] of arrangement patterns while fixing the colony geometry, employing a box counting algorithm for this purpose (Fig. 1E and Supplementary Sec. S.6). We observed that for the case of cell arrangement patterns formed by partitioning of colony into progeny chains, the Shannon entropy values is much lower than for the case of randomly intermixed patterns (Fig. 1F). This reinforces the ordered nature of the spatial distribution of cells in bacterial colonies. Indeed, the low values of Shannon entropy as well the decreasing propensity of entropy values even as the colony grows and self-organises (Fig. 2A), point towards the ubiquity of low entropy and entropy reduction in real life patterns and situations [32, 33], indeed a hallmark of life itself [34].

We next asked whether the cells in the two enclaves displayed a bias in the growth prospects towards cells in one of the progeny chains, due to the “vagaries of topology” as the colony geometry evolves. To do this, we measured the difference in phenotypic traits for cells belonging to the two domains. Firstly, we observed that the spatial areas of the domains are approximately the same, which is around half of the colony area (Fig. 1G), which is striking as the area comprises of cells as well as intercellular voids, pointing to a similarity in the way of cells accommodate and adjust spatially in the two enclaves. Further, mean cellular length, mean elongation rate of cells and the average area of cells in the two domains show very close similarity with the colony level statistics of the same (Fig. 1H, Supplementary Fig. 7 and Supplementary Fig. 8, respectively). A critical feature of bacterial cells, from the standpoint of colony survival and propagation, is their division into daughter cells. To compare and contrast colony level division statistics with that of the two domains, we calculated the number of division events occurring in the colony as well as the domains as a function of time. While at the single cell level, this can be a highly stochastic event (Supplementary Fig. 17A), we observed that as time progresses, even in this case, the colony level statistics show close proximity to the statistics of the two domains (Supplementary Fig. 9). This reinforces the fact that the two domains are a self-similar partition of the colony which transcend the dynamic nature of such partitioning. Further, we observed that in all cases, the two domains retained significant exposure to the surroundings of the colony and no “encircled” enclaves (in other words, sub-enclaves of cells from one progeny chain completely surrounded by cells from the other progeny chain) were formed as the colony grew. Indeed, the enclave perimeter exposed to the outer surrounding of the colony is around half of the colony perimeter for both the enclaves, even as the colony grows in size (Supplementary Fig. 10). This is important since boundary exposure is a crucial marker of survival fitness as it assures proximity to nutrients and space for expansion [13]. Thus, we conclude that in spite of growing freely in two dimensions and displaying a dynamically evolving colony geometry, the two domains originating from the progeny chains of the two initial daughter cells gain very equitable access to growth resources and resultantly, display very similar features as the colony at large.

## Activity drives dynamic partitioning of a colony into enclaves

One of the most noticeable features of bacterial growth is the dependence of biological activity on a range of factors, in particular ambient temperature, with cells growing and dividing faster at optimal temperatures. This difference in growth rates is quite apparent at colony scale but the effect of activity on the spatial geometry of the colony is completely obscured in standard imaging of cells, highlighting the importance of label-free tracking of cells in such colonies. We observed a somewhat counter-intuitive effect of activity on the spatial geometry of colonyat slower growth rates, cells of the two progeny chains displayed a more intermixed structure, while at higher growth rates, the colony is seemingly more ordered and less intermixed. The time series of Shannon entropy values show that slower growing colonies consistently display higher entropy of cellular arrangement patterns (Fig. 2A). Similarly as time progresses, the distance between the centroids of the two enclaves gets lower and proportion of inter-enclave contacts is higher for slower growing colonies (Fig. 1D and 2B). All this confirm the higher degree of intermixing and less ordered feature of colonies of cells growing slowly. We also observed a marked preference for narrow front invasion in this case, with thin fingers of cells invading into the region of the other, while on the other hand in case of faster growing cells, wide front invasion in seen with the invasion front comprising of many cells. To quantify this difference, it is necessary to delve deeper into the geometry of interfacial region of the two domains (Supplementary Sec. S.7). For this, we first computed the interfacial length normalised by the perimeter of the colony. In this case, we see that the interfacial length is much longer for the case of slow growing colonies (Fig. 2C). Further, high curvature regions along the interfacial curve are more numerous in case of slowly growing colonies compared to faster growing colonies (Fig. 2D-G). Presence of high curvature regions attest to the meandering nature of interfacial curve in this case as well the affinity for narrow front invasion, which is characterised by sharp turns and consequently, high curvature hotspots in the interface curve (Fig. 2D,E), an analogy of which can be drawn between river streams in hilly regions, characterised by tight bends and meandering curves compared to straight wide river streams in the plains. We also quantified the size of invasion front by computing the mean invasion width (Supplementary Sec. S.7), which necessitated a precise identification of the invasion areas in the interfacial region (Fig. 2D and Supplementary Fig. 11). Again, we observed that, compared to the case of colonies of fast growing cells, the mean invasion width is much smaller in case of colonies of slow growing cells, highlighting the preference for narrow front invasion in this case (Fig. 2H). Finally, to understand the nature of invasion front in the vicinity of high curvature regions, we calculated the probability distribution of invasion width in high curvature regions, observing that for slower growing colonies, peak is attained for lower values of width while faster growing colonies display a higher peak value of invasion width (Fig. 2I), which reiterate their propensity for narrow front invasion and wide front invasion, respectively. Therefore, we conclude that biological activity modulates the emergent spatial geometry of intermixing of cells in colonies, with higher activity resulting in more ordered, less intermixed colonies while lower activity promoting higher degree of intermixing, characterised by narrow front invasion of fingers of cells into the territory of the complementary enclave. It is notable, at initial times after just a couple of division events, the colonies all displayed similar arrangement of cells in all cases and evolving with time into disparate arrangement patterns (Supplementary Fig. 12), depicting that the dependence on activity level of cells on the arrangement patterns acts out after the colony has grown beyond a threshold size, reflecting the intermediate scale at which the phenomena takes effect. This is further highlighted by the feature of such colonies to self organise into a tapestry of microdomains of small numbers of similarly aligned cells, but in which case the size of such microdomains decreases with increasing activity [10]. Interestingly, *V.cholerae* cells which were grown at ∼ 25 *^°^*C also showed remarkable similarity of values of Shannon entropy, centroid displacement and normalised interfacial length with colonies of *E.coli* cells (Supplementary Fig. 13B-D), which corroborates that the dependence of spatial arrangement of cells on temperature mediated activity is robust across cell shape and species.

## Genealogical interface as a hotspot of orientational disorder

Next, we sought to understand the mechanics of enclave invasion and its effect on orientational order of cells. Local orientational order of cells in nonmotile bacterial species growing in colonies has been shown to emerge from the interaction of steric forces of cells shoving each other for space on one hand and active extensile stresses due to cell growth on the other hand, with the colony thus behaving like an active nematic liquid crystal [10, 11]. We hypothesized that enclave invasion will necessarily involve a high degree of jostling of cells, with the interfacial region likely emerging as a hotspot of orientational disorder. To investigate this, we compute the orientation order parameter *S*, a metric that encapsulates the local order in aligned systems, with *S* = 1 denoting perfect alignment while low values of *S* signifying high level of disorder in cell alignment [35]. For this, we first determined the local orientation which was appropriately averaged spatially to extract the value of *S* (Supplementary Sec. S.8). We observe that regions of low values of *S* in the colony tend to be along its boundary and along the enclave interface (Fig. 3A and 3B). Indeed, if we restrict ourselves to the interior of the colony, such regions of orientational disorder occur almost exclusively along the enclave boundary (Fig. 3B and Fig. 3C). We next tracked the location of topological defects in the colony *vis-a-vis* the spatiotemporal evolution of the enclave interface (Supplementary Sec. S.8). Topological defects, which are singularities in local orientational field, are associated with a breakdown of local orientational order and thus, regions of low values of *S* are markers of the presence of defects [14]. In case of bacterial colonies, only +1*/*2 and −1*/*2 defects were observed, as is well known [11, 15]. In our case, we observed that topological defects of both +1*/*2 and −1*/*2 charge proliferate in the vicinity of the enclave interface (Fig. 3B). Grouping defects into those that are in the vicinity of the interface, those lying near the boundary and those lying elsewhere, we observed that most defects are covered in the first two categories itself (Fig. 3C) and further, defects in the interior regions of the colony almost always arise in the vicinity of the interface (Supplementary Fig. 14C and Supplementary Fig. 15). Thus, while regions of high orientational disorder and presence of defects in the interior of bacterial colonies may seem random, label-free tracking of cells allows us unravel this riddle by exposing the active-active interface [36] of progeny enclaves as a hotspot for orientational disorder and topological defects.

**Figure 3:**
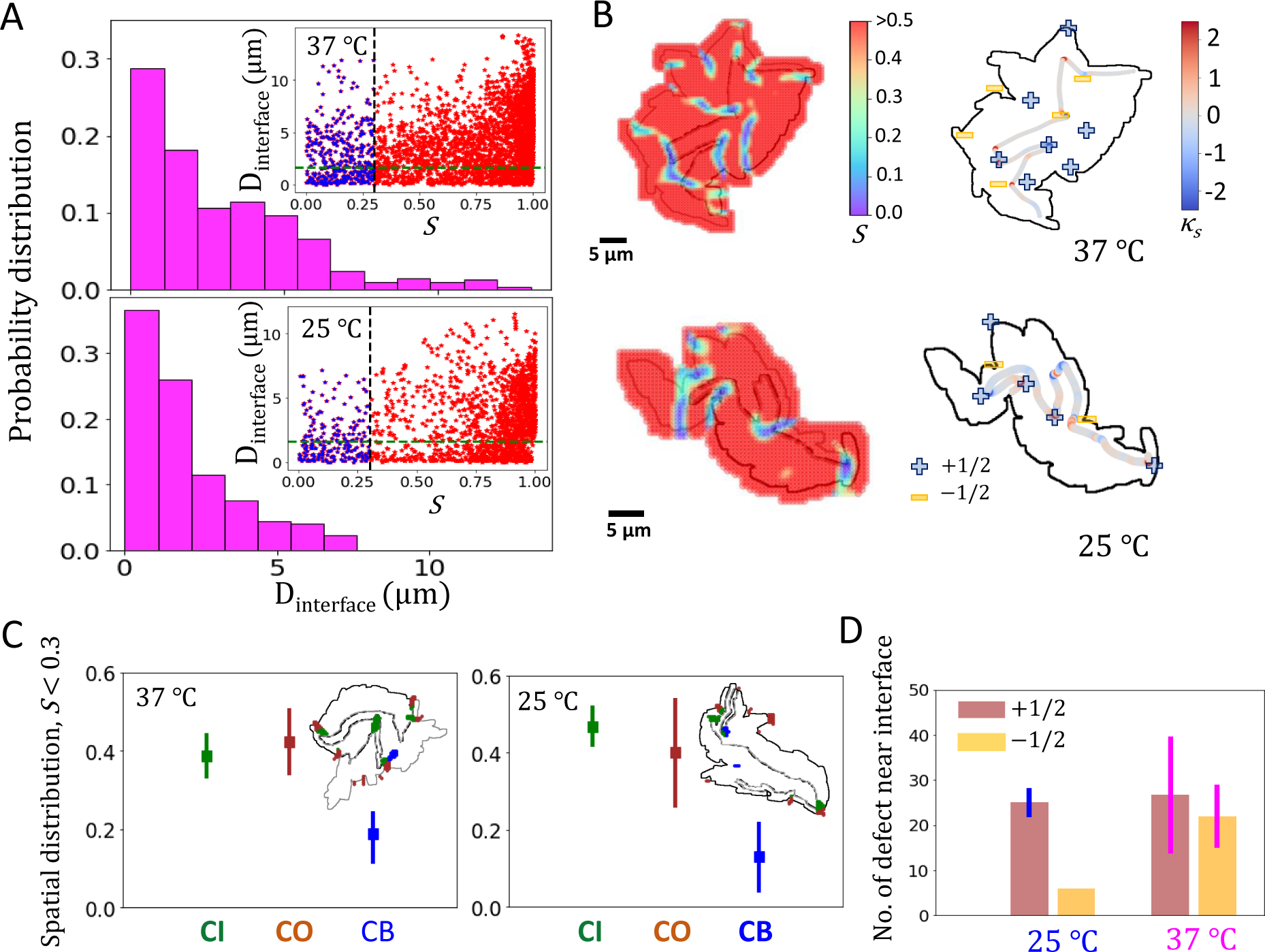
Proliferation of orientational disorder in the vicinity of enclave interface and its role in driving activity dependent enclave invasion. **A.** Probability distribution of low nematic order parameter (*S <* 0.3) regions as a function of distance from the interface for colonies growing at 37 *^°^*C (left) and 25 *^°^*C (right). Inset For a representative colony growing at 37 *^°^*C (resp. 25 *^°^*C), scatter plot of distance of grid points from the enclave interface is plotted as a function of nematic order parameter. The vertical dotted line shows chosen threshold value of low order parameter (*S* ∼ 0.3) and the horizontal dotted line shows the distance (*d_i_*) from the interface where most of the low order parameter points are lying (*d_i_* ∼ 1.5 *µ*m). **B.** Colormap of distribution of values of nematic order parameter *S* in shown for a representative case for colony at 37 *^°^*C (top left) and 25 *^°^*C (bottom left) and corresponding distribution of +1*/*2 and −1*/*2 defects are shown for the two cases (37 *^°^*C (top right) and 25 *^°^*C (bottom right)). **C.** Spatial distribution of low S points (plausible sites for topological defects) that are located in the vicinity of enclave interface (denoted CI), near colony boundary (denoted CO) and in the rest of colony bulk region (denoted CB) for colonies growing at 37 *^°^*C (left) and 25 *^°^*C (right). **D.** Distribution of +1*/*2 and −1*/*2 defects in the vicinity of enclave interface is shown for colonies growing at 25 *^°^*C and 37 *^°^*C.

We aimed to further localise regions of high orientational disorder and defects *vis-a-vis* the enclave interface. Firstly, we note that the majority of the cells in the interfacial region preferentially align parallel to the enclave interface (Supplementary Fig. 14A), as has been observed for interfacial regions in several other contexts [36, 37, 38]. Nevertheless, this preference is broken in regions of high curvature of the interface curve and we observed that regions of high disorder in the orientational field as well as presence of topological defects correlated strongly with regions of high curvature along the interface curve (Fig. 3B).

We also tracked the dynamics of defects *vis-a-vis* enclave invasion and observed that enclave invasion is typically initiated by topological defects as one enclave pushes into the region of the other. Over time such invasions developed while the initial defect persisted in the region, signifying the role of defects in nucleating and fostering the various modes of invasion (Supplementary Fig. 16). This is highlighted by looking at the distribution of defect type in enclave interface vicinity as a function of activity (Fig. 3D) which shows that −1*/*2 defects manifest in much higher numbers for faster growing cells. In fact, the geometry of the defects seem closely related to the invasion mode fostered by such defects. Defects of charge +1*/*2 have a comet like shape and behave like self propelled particles [39]. This, and the fact that they travel comparatively larger distances on average (Supplementary Fig. 14), points to their typical tendency to form and develop narrow front invasions which penetrate deeper (Supplementary Fig. 16). On the other hand, −1*/*2 defects have a tri-fold symmetry and in fact, display a propensity to be “caged” in the sense they move comparatively shorter distances, showing a saturating trend in their movement (Supplementary Fig. 14E). This attributes their tendency to nucleate and develop wide front invasion, which are hampered by their symmetry from penetrating deeper but rather slowly spread over a larger region (Supplementary Fig. 16). Indeed, this contrasting feature of the movement of +1*/*2 and −1*/*2 defects has been noted before, for bacterial colonies [11] and in case of other biological aggregations as well [40].

## Lattice model simulations capture the evolution of genealogical enclaves

To understand the physical principles underpinning the enclave formation in bacterial colonies, we employed a coarse-grained lattice cell model. We started with the two initial cells following the first division event placed next to each other, which were then progressively allowed to divide. The division time for each cell was chosen randomly following log normal distribution with the parameters of the distribution determined from our biological data (Supplementary Fig. 17A), with number of growth of cells in the colony and in the two enclaves showing good agreement with the experimental data, staying nearly equal with time (Fig. 4D and Supplementary Fig. 17C). To approximate daughter cell placement after cell division in colonies, we identified the eight sites nearest to the each site (referred to as the “Moore” neighbourhood for each site) and upon division, one daughter cell was allowed to maintain the position of its mother cell while the other one was placed in one of the neighbouring sites that was unoccupied, with equal probability (Fig. 4A). To approximate cell jostling in limited space in colonies we introduced cell shoving algorithms, for division of cells whose Moore neighbourhood becomes completely occupied - the shoving direction is chosen according to a probability distribution which ensures that cells will prefer to shove in directions where they have to move less number of cells (Fig. 4B and Supplementary Sec. S.9, we also simulated a shoving scheme where the shoving direction was chosen at random with equal probability, obtaining similar results, Supplementary Sec. S.9 for more details). The cell agglomerations were then simulated to grow till they reach sizes consistent with our biological data. We observe that in all cases, enclave formation was well displayed by the simulated colonies, with very few isolated and encircled progeny domains and the two domains displaying a very similar size to each other as they grow (Fig. 4C-E), showing a qualitative agreement with experimental data.

**Figure 4:**
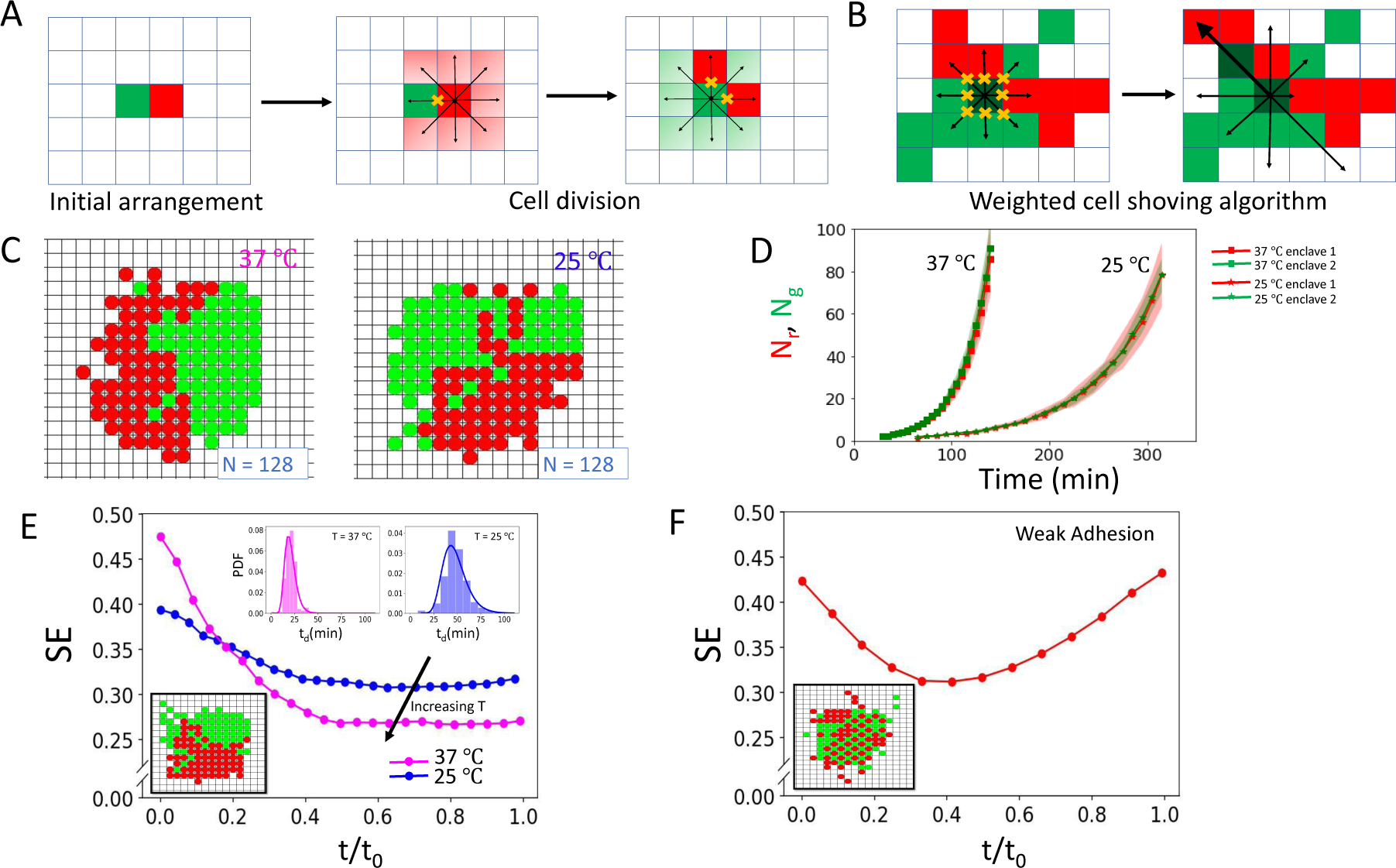
Lattice model simulations of bacterial colonies showing formation of enclaves in qualitative agreement with experiments. **A.** The simulation starts with two cells placed next to each other randomly. Division time for cells are assigned according to a log normal distribution whose parameters are determined from our biological data. Once the cells divide, one of the daughter cells occupy the original site while the next is assigned an empty site in its Moore neighbourhood, assigned randomly with equal probability. **B.** Once the Moore neighbourhood is completely occupied, upon cell division, one of the daughter cell occupies the original site and the other “shoves” the cells in a direction chosen according to a probability distribution so as to ensure that cells prefer to shove in directions which have less number of cells that must move. **C.** Lattice model simulations of colonies growing at 25 *^°^*C and 37C are shown here for two representative cases with number of cells N=128 and the two progeny chains colored red and green. **D.** The number of cells in each domain is plotted as function of time. **E.** Shannon entropy of arrangement pattern is plotted as a function of normalised time for lattice model simulations with high (colored magenta) and low width (colored blue) division time statistics to simulate colonies growing at 25 *^°^*C and 37 *^°^*C, respectively. (Inset-) Distribution of division times and snapshot of a simulated colony. **F.**Evolution of Shannon entropy of arrangement pattern with normalised time for the case where a daughter cell is allowed to be placed away from neighbourhood of the mother cell (in particular two lattice steps away), thus not requiring two daughter cells to always be neighbours, in effect weakening the adhesion of daughter cells, (Inset-) Snapshot of a simulated colony in this case.

We interpret this observation as mixing-demixing in binary mixtures, where the free energy of the system *F* is given as *F* = *E_i_* −*TS^mix^* [41, 42, 43], with *E_i_* being the energy of interaction and *S^mix^* the mixing entropy, which promotes mixing of the components. In our case, intermixing is driven up by the shoving of cells in the colony as they jostle for space, which drives them farther from their genealogical milieu and promotes mixing of progeny chains. On the other hand, as we can observe from colony growth images, cells remain in contact after division. This associates an energy cost to cell shoving, for instance to separate a cell from its sister, to effect mixing. To test this, we simulated the formation of colonies in which, upon division, cells were allowed to occupy sites not just in the neighbourhood of the mother cell but a step farther in the lattice as well (Supplementary Sec. S.9), thus in effect weakening the energy favouring interaction of cells from their own progeny chain. We observe that such a minor change is in fact sufficient to cause distinctly different behaviour, in particular exhibiting a high degree of intermixing and resulting in an increase in Shannon entropy of the arrangement pattern with time (Fig. 4F, Supplementary Fig. 18C,D). Thus, while viable colonies are still formed when the cells are allowed to separate upon division, spatial demixing of progeny chains weakened and a high degree of intermixing became apparent. This underscores the importance of adhesion of daughter cells upon division, likely due to self recognising adhesion molecules or mediated by EPS production [44, 45], for the formation of genealogical enclaves.

Next, we wanted to probe the effect of self similar growth of progeny chains, as displayed by bacterial colonies as well as our simulations (Fig. 1G and Fig. 4D), in promoting enclave formation. For this, we differentiated the division times of cells by slowing down the division time for cells belonging to one progeny chain. We see that for small differences in division time, enclave formation is still prevalent (Supplementary Fig. 19). This suggests that formation of progeny enclaves in microbial colonies can remain robust even when varying growth levels of cells results in difference in the number of cells in the progeny chains as they grow (for instance when wild type cells and mutant cells having a small variation in growth rates are grown together). However, when the difference in growth rate is large, this leads to large imbalance between the number of cells in the progeny chains, particularly as the colony grows large and we observe a high order of intermixing between the cells of the two progeny chains (Supplementary Fig. 19). Notably, this is similar the case of binary mixtures in which case, for low concentrations of one of the components, mixing is displayed as demixing becomes energetically prohibitive [43].

Finally, we note that our simulations also showed a similar dependence of intermixing on activity as our analysis of bacterial colonies had established. Specifically, we calculated the proportion of inter-enclave contacts of cells in the colony and the Shannon entropy of the cell arrangement patterns obtained from the lattice model simulations and observed that both the inter-enclave contacts and Shannon entropy values are consistently lower for the case of faster growing colonies (Fig. 4E and Supplementary Fig. 17E,F and compare with Fig. 2A,B), which suggest a lower degree of intermixing in this case, as was the case for bacterial colonies. Since activity here is encoded by the division times gleaned from our biological data with more active colonies have a wider spread of division times, thus the lattice model simulations suggest that stochasticity in cell division times, engineered by biological activity, is a factor in determining the disorder in the emergent organisation of cells in bacterial colonies. This is likely due to effective space grabbing by both progeny chains for faster growing cells, whereby the lesser spread in division times ensures that space is filled in orderly fashion with both progeny chains occupying sites one after other. On the other hand, the wide spread in division times can sometimes lead to cells from one progeny chain dividing more than once before a cell from the other chain can divide, leading to differential site occupation, locally in space and time. However, the large difference in Shannon entropy values for cells in bacterial colonies suggests that there are other factors at play as well. The characteristic patterns forming due to enclave invasion are reminiscent of mixing of viscous fluids[46, 47], which typically occurs when a low viscosity liquid displaces a high viscosity liquid under pressure. In case of enclave invasion, while the effective viscosity of the enclaves will remain the same, activity drives the invasion. Further, for cells growing faster, it is likely that the cell-cell tensions are higher in the interfacial region [48], which can increase effective viscosity for cell aggregates, resulting in blockage of deeper thin invasions and giving rise to wide, shallow invasion fronts with more ordered cell arrangement patterns as observed.

## Conclusions

To summarise, we have exhibited the emergent formation of genealogical enclaves in growing bacterial colonies by leveraging label-free tracking algorithm to trace the spatiotemporal evolution of progeny chains emanating from the first two daughter cells of the colony founder. The formation of such progeny enclaves was observed even for colonies of cells having low aspect ratio and even when substrate properties were changed. These enclaves are self-similar to the colony at large on several key phenotypic traits, thus showing the equitable growth prospects of the progeny chains and defying factors like jostling of cells for space and stochasticity in division times which could have potentially obstructed it. A key determinant of microbial life in their activity and while it is easy to discern the growth rate of colonies to differentiate between faster and slower growing cells, unravelling the effect of activity on spatial organisation of cells in such colonies is not straightforward. However, progeny tracking allowed us to highlight important differences in the way cells arrange themselves in colonies as a function of activity. We showed that the dynamic partitioning of cells of the colony in enclaves is activity dependent. While slow growing cells tended to intermix more amongst the two progeny chains and displayed a preference for narrow front enclave invasion, faster growing cells displayed less intermixing and a preference for wide front enclave invasion. We also demonstrated that the enclave interface is a hotspot for orientational disorder with regions of high disorder and presence of topological defects in the colony interior typically manifesting in the vicinity of it. Activity dependence can be discerned in this case as well, with faster growing colonies often displaying a relatively high proliferation of −1*/*2 defects in interface vicinity, whose tri-fold symmetry and sluggish mobility correlates with the typical wide front enclave invasion observed in this case. This also further highlights the utility of progeny tracking whereby seemingly randomly manifesting regions of high disorder in the colony in fact commonly turn up at the interface of genealogical enclaves. Finally, we employed a lattice model to understand the robustness of enclave formation, showing that the propensity of daughter cells to “stick” to each other upon division underpins the formation of genealogical enclaves and the characteristic “far from random” cell arrangement patterns were observed in bacterial colonies. Drawing analogy from the thermodynamics of binary mixtures and phase separation, our simulations showed how even a minor weakening of the tendency of daughter cells to adhere after division was sufficient to engender lineage mixing, even as viable colonies may still be formed under such conditions. This also allowed to show that one of the ways in which activity can influence spatial organisation of cells in colonies is by the inherent level of stochasticity in the division times of cells in such colonies. Thus, our work shows that while it might be expected that the freely growing bacterial colonies will display a thoroughly intermixed arrangement geometry of cells, in fact the opposite holds, with segregation of cells belonging to different progeny chains being observed under a range of conditions. As is commonly known and observed even for human beings, the most meaningful interactions often take place between spatial neighbours on one hand and lineage relatives on the other hand. Our work demonstrates that for bacterial colonies, in fact spatial neighbours are commonly lineage neighbours as well, thus correlating spatial and genealogical proximity. Ultimately, this points to the importance of living in the vicinity of close kith and kin to maximise survivability and liveability.

## Acknowledgments

We gratefully acknowledge the support from the Institute for Advanced Studies, University of Luxembourg (AUDACITY Grant: IAS-21/CAMEOS to A.S.) and a Human Frontier Science Program Cross Disciplinary Fellowship (LT 00230/2021-C to G.R.). We thank J. Nguyen for the raw image displayed in Supplementary Figure 6 (A). A.S. thanks Luxembourg National Research Fund for the ATTRACT Investigator Grant (A17/MS/ 11572821/MBRACE) and a CORE Grant (C19/MS/13719464/TOPOFLUME/Sengupta) for supporting this work.

## Author contributions

Conceptualization, planning, administration, and supervision: A.S. Methodology: A.S. and G.R. Investigation, data and statistical analysis, computations and modeling: G.R. with inputs from A.S. Writing: G.R. and A.S.

## S Supplementary Materials

### S.1 Bacterial cultures

The primary strain used in this work is a Gram negative bacteria *E.coli* strain, namely NCM3722 delta-motA. The second species considered in the study is *V.cholerae*, to understand the effect of cell shape (specifically low aspect ratio) on the spatial distribution of progeny chains. The cells were first streaked on a standard lysogeny broth (LB)-agar plate and grown by setting incubator at appropriate temperature. After a day of growth in the plate, single isolated colonies were identified and picked using a inoculation loop and transferred to liquid LB media in a shaker and shaken at 170 rpm. The cells were allowed to grow and divide in the shaker till late exponential phase (with regular subsampling done to track the cell growth over time by measuring the optical density (OD)), after which the cells were transferred to fresh LB media at a dilution factor of 1:1000 (ratio of bacterial suspension to medium) and grown for ∼ 1.5 − 2 hr (early exponential phase). Around ∼ 1 − 1.5*µ*L droplets were then inoculated onto a thin layer specially designed substrate. At least three biological replicates for each case were considered in this study. More details of the experimental protocol may be found in Ref. [1].

### S.2 Agar pad preparation and Time lapse Imaging

Low melting agarose was mixed with LB medium and the gel-like solution was poured into the Gene-frame (Thermo Scientific Gene Frame with thickness ≈ 0.25 mm) pasted on a standard microscopic glass slide and with a cover-slip to seal from the top. The gel concentration for most of the studies presented in this work is 1.5%. The effect of substrate properties was incorporated by varying the gel concentration in the range 1% to 2%. Growing colonies were observed in phase contrast mode using Olympus IX83 microscope with a 60 X oil objective (Hamamatsu ORCA-Flash camera). The whole system was set inside a thermally insulated temperature-controlled incubator (Pecon). First we located the coordinates of isolated cells on the agarose pad and the microscope was automated to capture these pre-recorded coordinates and record the images of the colonies as they evolved (for the case merging of two colonies, two nearby cells were located and images were recorded as cells grew and divided forming colonies that eventually merged). The images are taken at regular intervals (typically 2 − 3 min). Analysis of phase contrast images was performed using the open-source softwares Fiji:ImageJ [2] and Ilastik [3] and custom written Python codes.

### S.3 Image segmentation and label-free tracking

The phase contrast raw images were pre-processed by adjusting brightness and subtracting background noise using ImageJ. Then further background filling was done by applying tophat and black-hat filter using Python-OpenCV. A number of these pre-processed images were trained in Ilatik by pixel classification of cells and background. The trained classifier was then applied to rest of the frames by using batch processing in Ilastik. This training step involved several iteration until high order of segmentation was achieved. The segmentation was done for frames until the colony attained MTMT which was carefully noted manually by checking the pixel intensity of the images (Fig. 1A-D). For the segmented cells, length (denoted *l_c_*) was obtained by calculating the distance between the cell centroid and the center of two poles (which are two extreme ends of the contour of a cell). Cell width (*w_c_*) was obtained using length and area of cell contour (*a_c_*) using 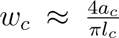. The length and width values thus obtained were further confirmed matching with extracted values of the major and minor axes length of each cell contour using the openCV ellipse fitting. The ellipse fitting also helped us to extract average orientation angle of cells. The colony outer boundary was extracted by image dilation and filling, which was used to extract the outer boundary perimeter and effective colony area.

Cell progeny chains were spatially mapped using a custom built tracking code written using python. To do that, first we extracted the features of the cells in all frames- centroid positions(*x_c_*,*y_c_*), length (*l_c_*), width(*w_c_*) and a label (*c_k_*) was assigned to all the cells. We compare two consecutive frames (here referred as frames *t* and *t* + 1) by looking for the “only” cell in frame *t* + 1, whose centroid is within a cutoff distance *d*(*c*) from the centroid of a cell in frame *t* (initial *t* = 0 is the time when the first pair of daughter cells are born). This pair of cells whose centroid displacement is within the cutoff were assigned to the same progeny chain. The cutoff value is appropriately chosen and optimised, being around the order of cell width. This covers most of the growing cells, but a few cells may be left out in frame *t* (either the cell in frame *t* has divided into two cells in frame *t* + 1 or have undergone substantial movement compared to the cutoff distance). To track the cell which divide in *t* + 1, we introduced “dummy” centroids for cells in frame *t*. Dummy centroids are points between the poles and original cell centroid, chosen by anticipating the centroids of the two daughter cells if the cell were to divide in the next frame. This helped us to map the dividing cell in frame t to daugther cells in frame t+1. A small number of cells may remain after this, which may have undergone substantial movement (especially cells located near colony boundary which have free space to move). These were mapped by noting the displacement of their poles and the centroids within a cutoff distance in the two consecutive frames. If necessary, manual correction was done to rectify identification errors. Thus, we can spatially and temporally track the progeny chains emanating from the first two daughter cells (Supplementary Fig. 1E,F and Supplementary Fig. 2).

### S.4 Cell contact number analysis

We compute the average cell contacts (neighbours) of cells in the colony. For each cell *C* in a given time frame, we first computed distance between centroid of cell *C* and centroid of other cells, and we collected the cells ({*C_r_*_0_}) which were within cutoff radius *r*_0_ (∼ 2 − 3 cell length) of cell *C*. For each cell in {*C_r_*_0_}, we first selected a large number of points on their boundary and calculated the distance from points on the boundary of the cell *C*. If the minimum distance between two cell boundaries was less than a small tolerance value (≤ *w_c_* width of the cell), then we called them neighbours ({*N_r_*_0_}). The neighbours were matched manually in several instances to ensure correctness. Following this, we then looked at the fraction of inter and intra contacts (at the level of progeny chains) based on the descendants of the first two daughter cells.

### S.5 Global properties of the enclaves

We now discuss how the centroid, perimeter, common interface length and area of enclaves formed by progeny chains of cells were calculated. First we extracted the arrangement of the cells in the two enclaves based on the cell label information we get from the tracking algorithm and thus, the outline of their outer boundaries. Then, we computed their centroid coordinates using using OpenCV (the calculations were done frame by frame).

To estimate the common interface length of the two sub-colonies, we first computed the perimeters of the outer boundary of the two enclaves formed by descendants cells, denoted *P_s_*_1_ and *P_s_*_2_. Since in all the cases the two enclaves are completely in the interior of the colony with common interface boundary, the interface length is simply given by

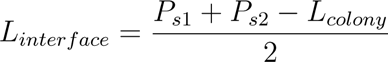

, where *L_colony_* is the perimeter of the of the entire colony. Again the calculation is carried out frame to frame to track their temporal evolution.

### S.6 Shannon entropy for cell arrangement patterns in colonies

We leverage the concept of Shannon entropy to compare and contrast disparate cell arrangement patterns in growing colonies, which we describe now. We employed a moving box algorithm (box size, *s* ≈ 5.5 *µ*m) with box moved progressively by distance *s/*2 to run through the cells in the colony. Cells of the two enclaves having been assigned colors green and red and represented by their centroids, we calculated the probability of seeing a red cell or green cell in each box. Thus, for every box *B*,

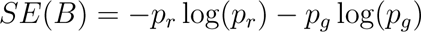

gives the Shannon entropy of the arrangement in the box itself, where *p_r_* and *p_g_* are probability of finding a red and green cell respectively. The average Shannon entropy for each colony was then calculated by averaging the entropy values obtained for each (moving) box. Next, to get a measure of the magnitude of the obtained entropy value in comparison to the entropy values obtained for all possible cell arrangement patterns, only fixing colony geometry and the precise number of cells in each domain, we assigned the two colors red and green in same proportion to cells in the colony at random and computed avg. Shannon entropy. This calculation was repeated in each case for 2 × 10^5^ iterations, to derive the range of values of entropy values sweeping through the phase space of all possible conformations and so as to estimate and compare the range of entropy values arising out of real life patterns.

### S.7 Geometric features of enclave interface

To quantify enclave invasion, we probed the curvature of the interfacial curve and other geometric properties of the invasion front. For computing the local curvature of the interface curve, we utilised the kappa plugin in Fiji [4]. Next, we calculated the area of the invasion of one enclave into other by a “hull” filling process to determine the region of invasion. Specifically, the high curvature regions which marked the beginning of the invasion front of one enclave into other were joined to demarcate a region of invasion, see Supplementary Fig.11. The mean invasion width for enclave invasion by calculated by measuring the width of the invasion fronts in several locations and taking their mean.

### S.8 Orientation field of cells and topological defect detection in the colony

For all phase-contrast images, we first measured the orientation by the structure tensor method. The structure tensor was computed by taking the outer product of the gradient vector with itself for each pixel, locally averaged within a given Gaussian window for a size chosen roughly one-quarter the size of single cell. To retrieve the local orientation stored in the structure tensor, eigenvalue analysis was performed. One of the eigenvectors of the structure tensor encodes the orientation value (Φ) at each pixel lying between −*π/*2 to +*π/*2 (Supplementary Fig.14A).

Next, we computed the nematic order parameter,

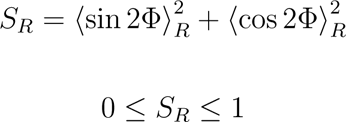

Here, the spatial average is done within fixed size square region *R* (with each square roughly containing 3 − 4 cells) and the moving grid algorithm with steps of atleast one-third size of grid size was employed to cover the entire colony for averaging (Supplementary Fig.14B). Only pixels that resided in the bacterial cells of the colony (white pixels) were taken into account to compute the local order parameter. This was done to get rid of the false orientations especially at the pixels near the cell boundary regions. Then we located the positions of low values of the order parameter by considering the points where *S_R_* ≤ 0.3. Of these points, local minima of the nematic order parameter values were taken as candidates for topological defect cores. For these possible defect sites, automatic defect detection was done by calculating the topological charge (*q*) [5]. Only two types of defects (charge +1*/*2 and −1*/*2) were found (Supplementary Fig.14C). Further manual correction of the defects is done by checking position of defects between atleast two consecutive frames.

### S.9 Lattice model of colony growth and enclave formation

We start with discrete square lattice with 25 × 25 sites, which is large enough to simulate colonies of the size that are of interest here. We assume that each lattice site can be occupied by a single bacteria and no site is allowed to overlap. The initial configuration is of a “red cell” that is placed at the center of the grid (daughter 1) and its sister a “green cell” (daughter 2) is randomly allotted a site in the Moore neighbourhood of the red cell. The simulation is then set off from this configuration, with the initial two cells dividing and their daughter cells dividing and so on and so forth, giving rise to their progeny chains. Every cell, starting from the initial two cells and then upon the birth of any new cell, is assigned a division time chosen at random using the log normal distribution, with the mean and standard deviation fixed to be the same as that gleaned from our biological data for colonies growing at 25 *^°^*C (48 ± 13 minutes) and 37 *^°^*C (21 ± 6 minutes), as shown in Supplementary Fig. 17A. Based on the assigned division times, at each timestep, we arranged the cells in ascending order, with the top most cell in the list selected for division (thus, in the simulation, each timestep corresponds to the time when a cell in the colony divides). Following each cell division event, we let one daughter cell remain at the same site as its mother cell site while the other daughter cell is allocated a randomly chosen site from the unoccupied Moore neighborhood of the site of the mother cell. The new cells are then assigned division times as described and the order of the cells according to division times is updated. These steps are repeated and new daughter cells continue to fill up the unoccupied Moore neighborhood of mother cells. As the number of cells increase, a stage is reached when the Moore neighbour of interior cells is completely occupied. For this, we introduce a shoving algorithm to allocate space for the daughter cells, whereby a direction from the mother cell site joining it to one of the sites in its Moore neighbourhood is chosen with the probability of that direction being picked being given a weight to ensure that directions for which fewer cells need to be shifted is preferred (thus, the weight for a direction is inversely proportional to the number of cells that lie in that direction that need to be shoved). The cells are shifted outward by one site in that direction (mathematically, this is referred to translation along the chosen direction), thus creating an unoccupied site in the vicinity of the mother cell where one of the daughter cells is then placed. This rule allows us to effectively mimic growing bacterial colonies where cells push and jostle with their neighbours to allocate space for themselves and their progeny (we also consider an alternative shoving algorithm where the direction is chosen at random with equal probability, however this does not change our results qualitatively. The simulation is continued in this manner till the number of cells increases to values consistent with those obtained from our biological data. Inter-enclave contacts of cells in these simulated colonies is calculated by looking at the number of cells lying in its Moore neighbourhood that belong to the opposite enclave. Shannon entropy is calculated by box-counting method, with box sizes chosen to ensure similar number of cells as in the case of Shannon entropy calculations as performed in Supplementary Sec. S.6, on our biological data. Averaging was done by performing over 20 simulations for each case (Supplementary Fig. 17E,F and Supplementary Fig. 18A,B). Further, we also consider the case where upon division cells are not placed in the immediate Moore neighbourhood but are also allowed to be placed in the Moore neighbourhood of Moore neighbourhood of the original cell (thus in the lattice lying two steps away), Supplementary Fig. 18(C) and (D). Finally, we also consider simulations where cells belonging to one progeny chain divide faster while cells belonging to the other progeny chain divide slower (Supplementary Fig. 19).

---

**Supplementary Figure 1:**
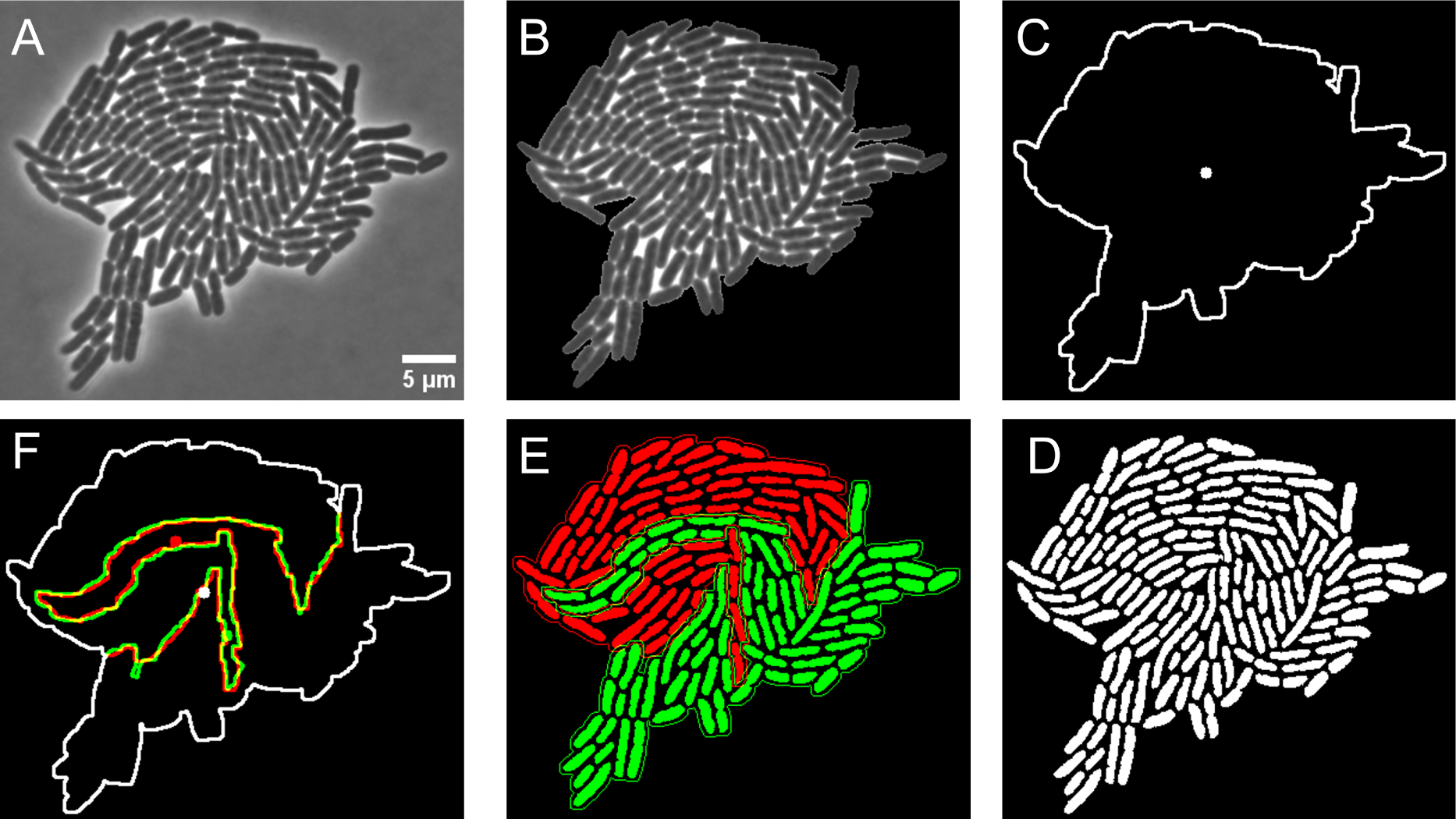
Image analysis pipeline. **A.** Phase contrasted raw image, **B.** Image contrast is adjusted and background filling done applying top hat and black hat filter using Python-OpenCV. **C.** Colony boundary and centroid (white dot) is extracted by deducing the outer contour using image dilation and space filling. **D.** Image segmentation is done by training in Ilastik. **E.** Label-free tracking was used to deduce the two progeny chains(red and green color denote the two enclaves formed by the descendant cells). **F.** Enclave centroid (red and green dots) and colony centroid (white dot) are shown within the colony boundary with the interface of the two enclaves shown by red-green overlapping lines.

**Supplementary Figure 2:**
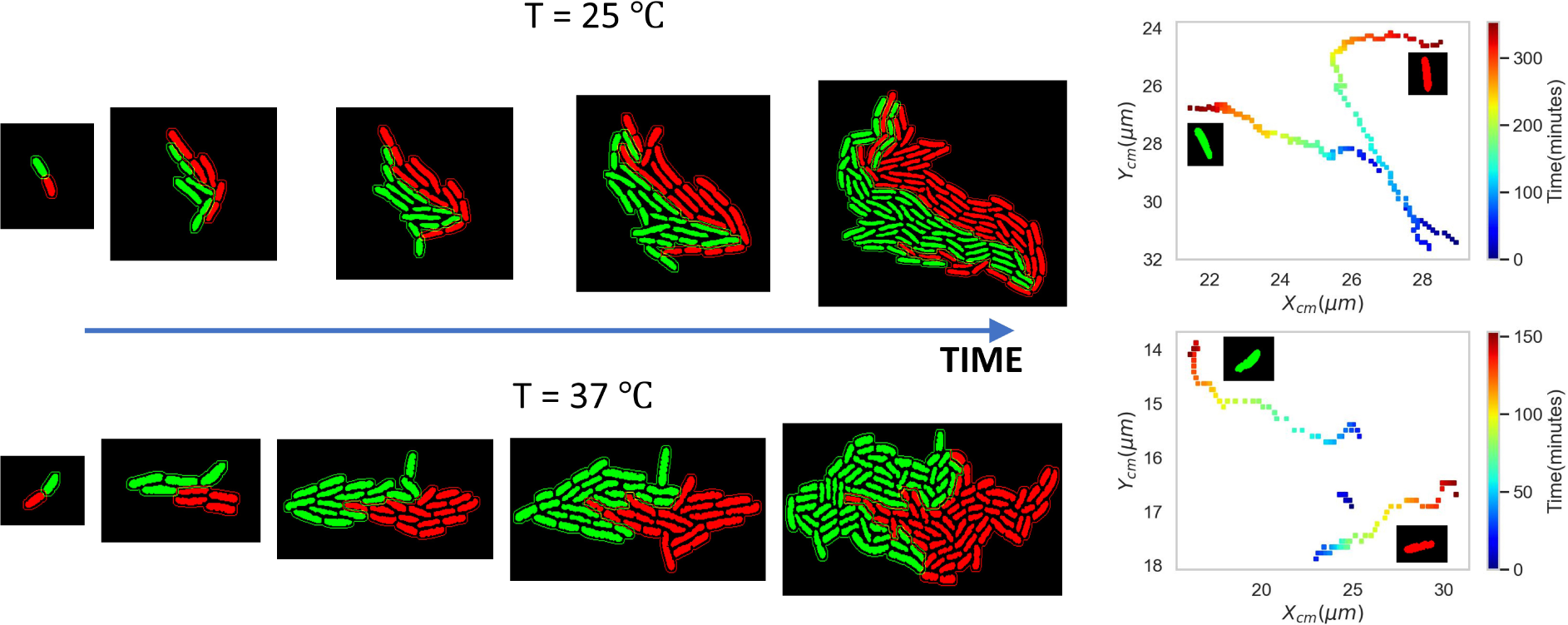
Enclave formation and evolution-. Spatiotemporal evolution of progeny enclaves is shown for representative colony growing at 25 *^°^*C (top) and 37 *^°^*C (bottom). On right, colormap of the position of the centroid of the two enclaves as they evolve with time is shown in each case.

**Supplementary Figure 3:**
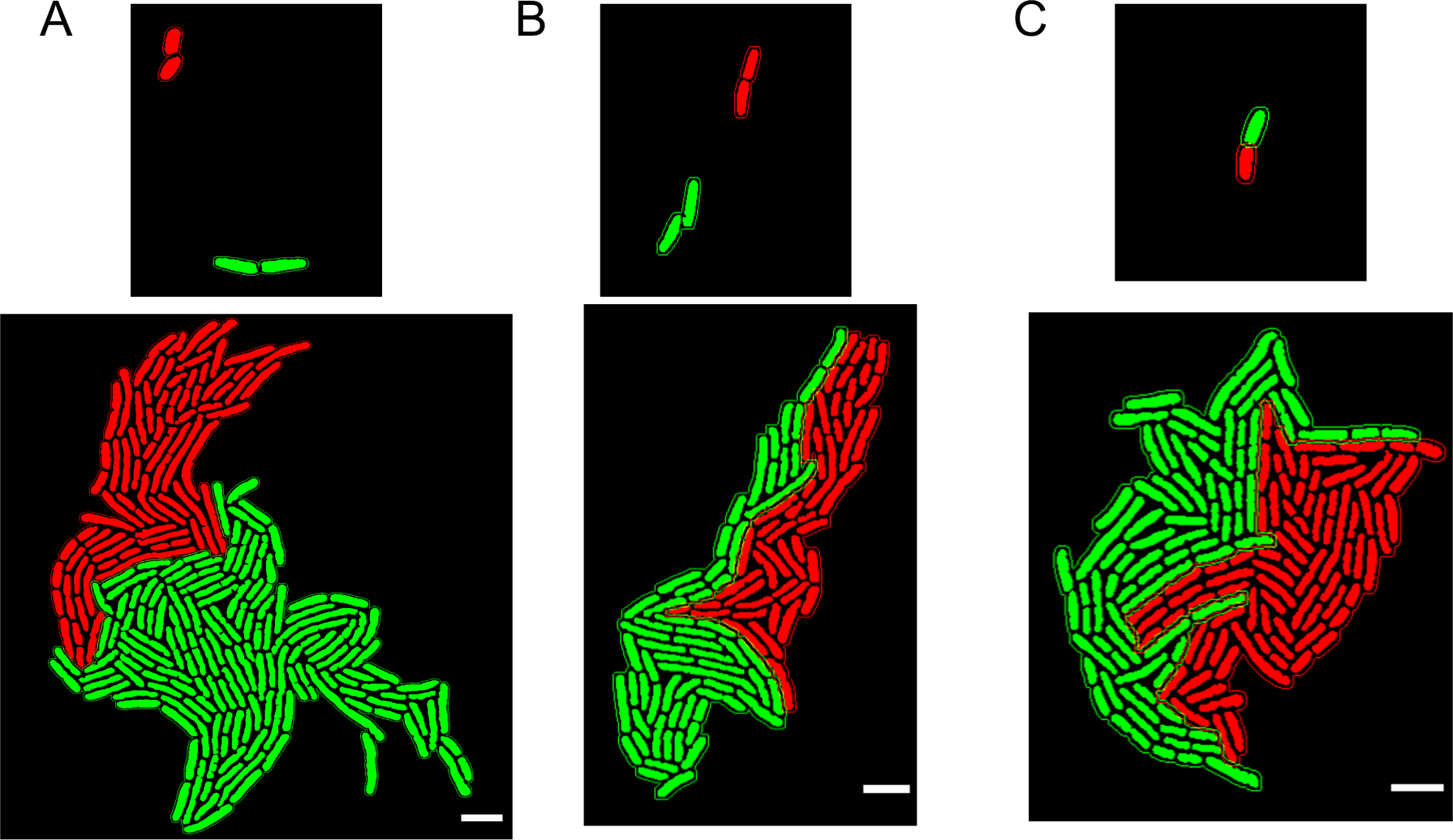
Cell arrangement patterns as two colonies merge and as the progeny enclaves emerge in single colonies are analogous. Starting with two founder cells, the merging of two colonies (colored red and green) to form a larger colony is shown here in two instances (**A** and **B**), with. Notably, the two colonies upon merging show very similar spatial organisation of cells in terms of partitioning of cells into enclaves as the case of progeny enclaves originating from a single founder cell, shown here for a representative case (**C**). The scale bar (white line) represents 5 *µ*m.

**Supplementary Figure 4:**
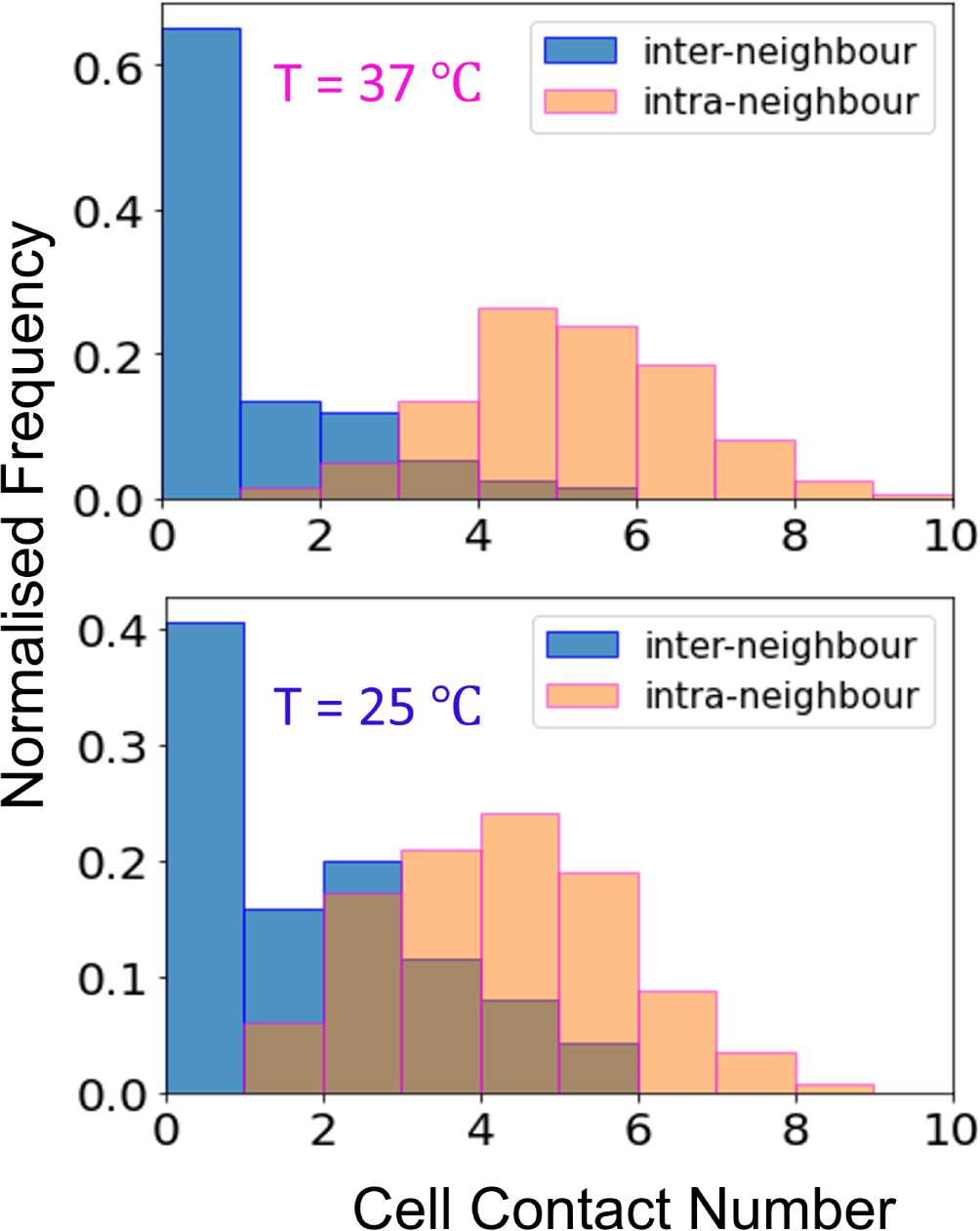
Cell contact number distribution-. Normalised frequency distribution of cell contacts with inter-enclave and intra-enclave contacts colored blue and orange, respectively.

**Supplementary Figure 5:**
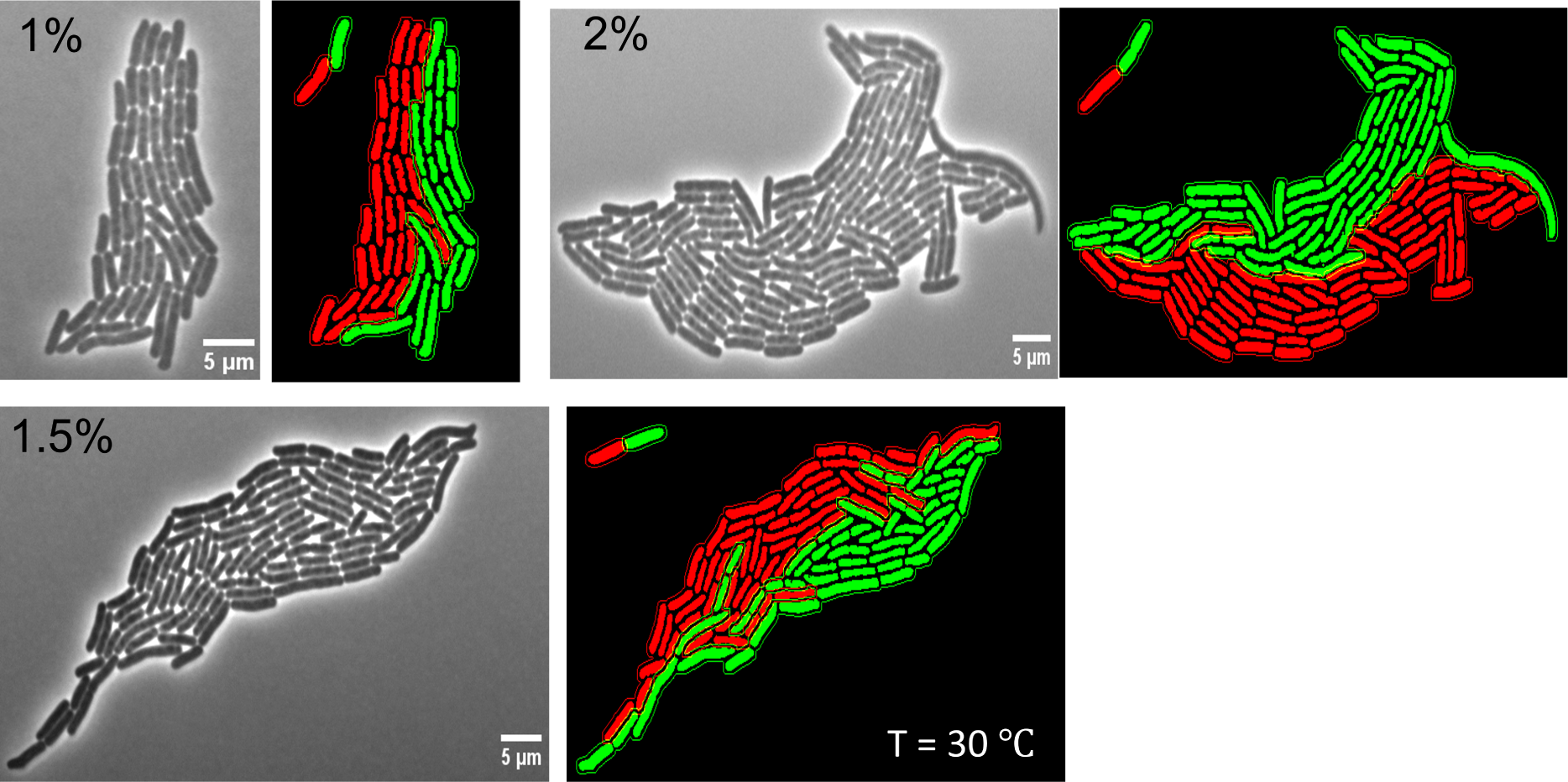
Colonies grown on varied substrates also show formation of progeny enclaves. Colonies grown on substrates with varying agarose concentration (1%, 1.5% and 2%) also display formation of progeny enclaves. The inset in all cases shows the two daughter cells arising from the first division event of the colony.

**Supplementary Figure 6:**
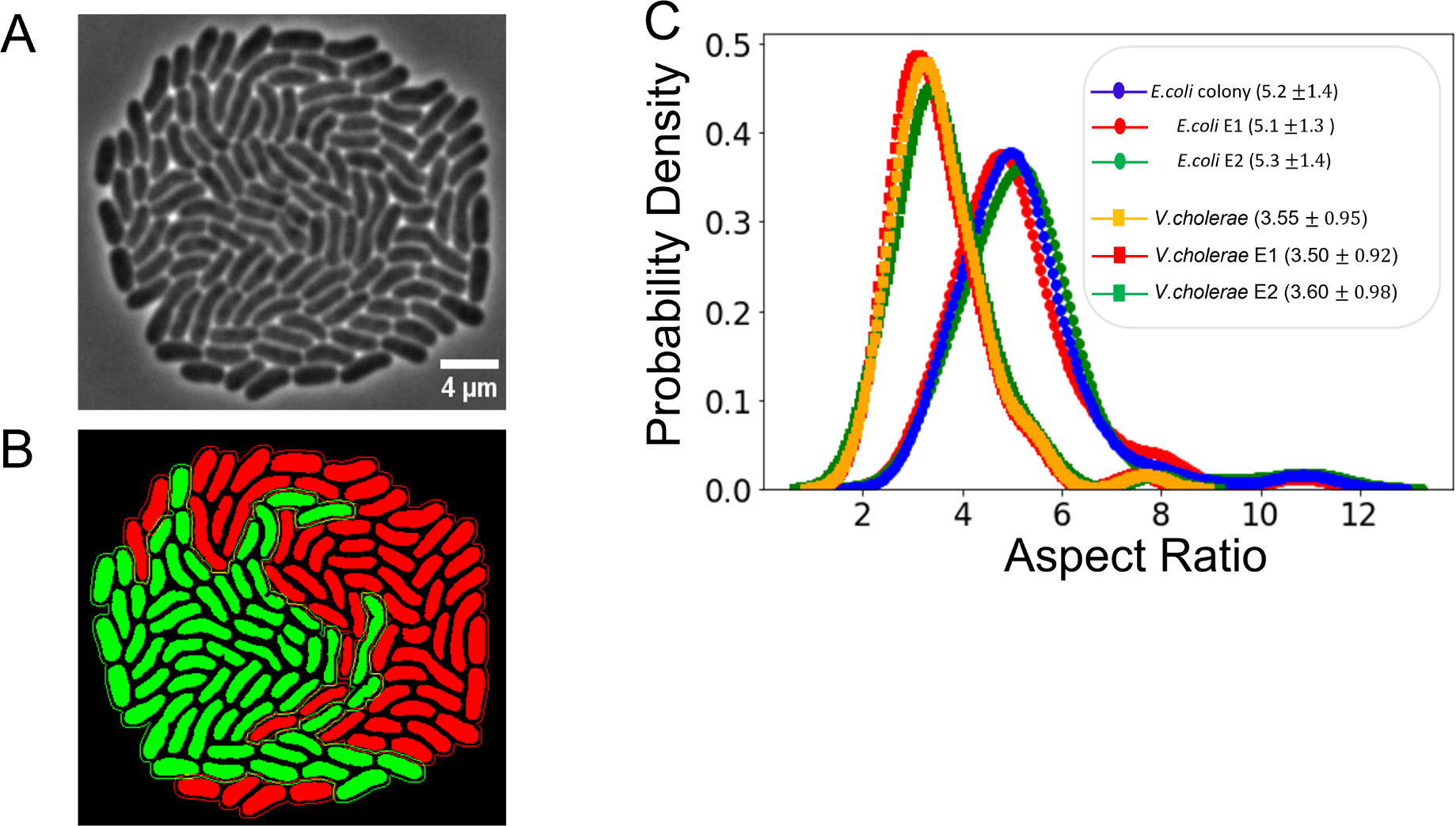
**Low aspect ratio cells also display enclave formation**. Progeny enclaves is displayed is case of other bacterial species as well, specifically *V.cholerae* growing at 25 *^°^*C. **A.** Raw image of the colony at MTMT (Courtesy of J. Nguyen), **B.** The appearance of enclaves obtained using label-free tracking suggests that it is robust for non-motile species of bacteria independent of cell shape. **C.** Probability density of the aspect ratios of the cells for *V. cholerae* and *E.coli* considered in this work, depicted for the two enclaves as well as the whole colony. The *V.cholerae* cells clearly have low aspect ratio (henceforth referred to as LAR strain). The probability density of the cells in the two enclaves show very similar aspect ratios in both cases, an exibition of self-similarity of the two enclaves for both the species.

**Supplementary Figure 7:**
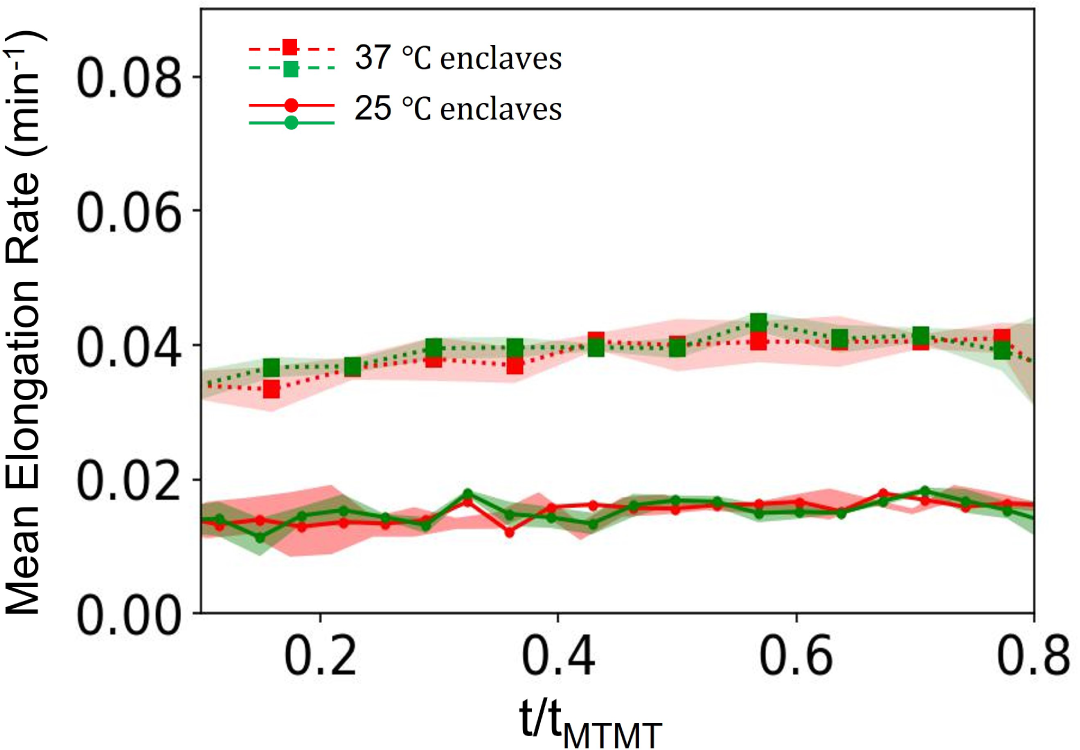
Self similarity between the enclaves and the colony as they grow. Mean elongation rate of cells in the two progeny chains (colored in red and green for both cases) is plotted as a function of time for cells growing at 37 *^°^*C (dotted lines with squares) and at 25 *^°^*C (lines with circles).

**Supplementary Figure 8:**
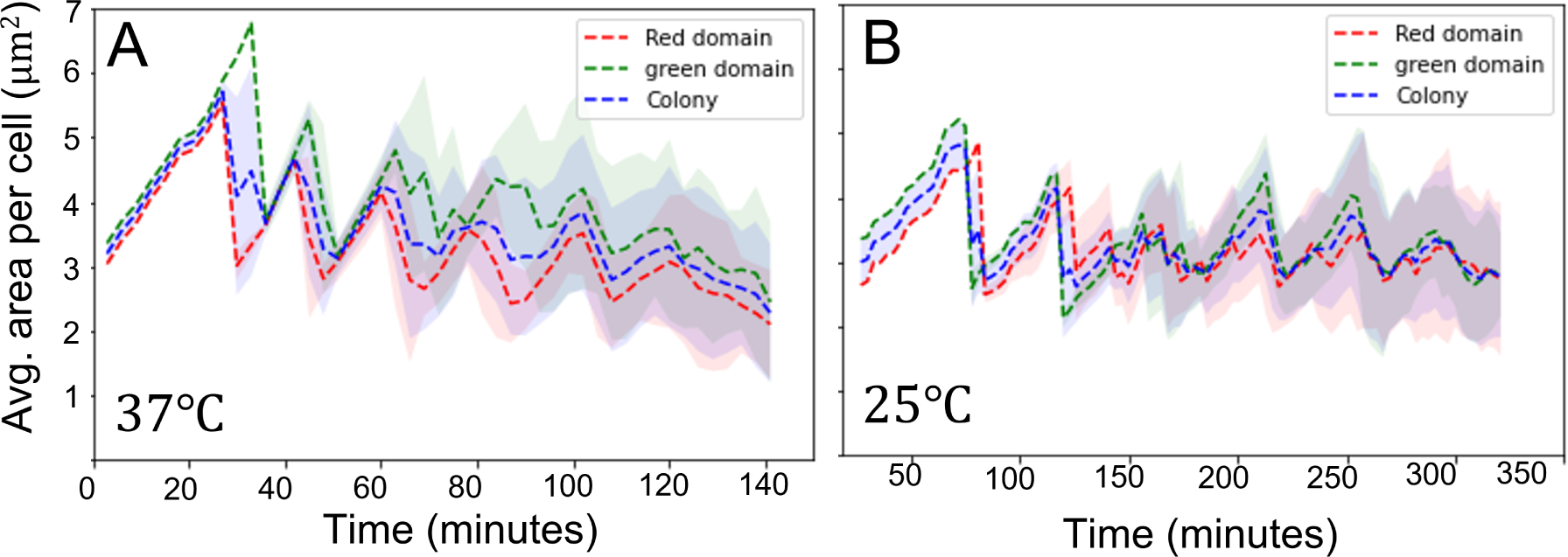
Self similarity between enclaves and colony in the average area occupied by a cell. Average area of a cell in the two enclaves (colored red and green) and the colony (colored blue) is plotted as a function of time for cells growing at 37 *^°^*C (**A**) and at 25 *^°^*C(**B**).

**Supplementary Figure 9:**
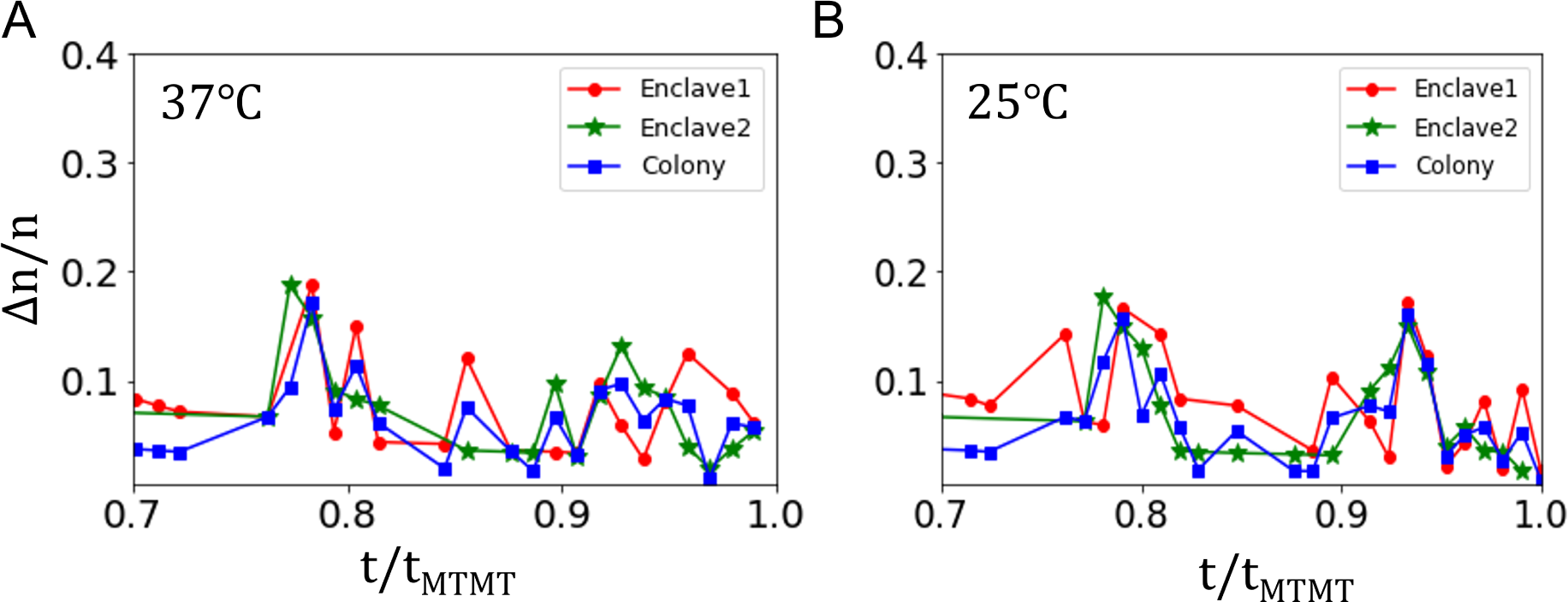
Division events in the colony and the enclaves evolve similarly with time. Normalised number of division events in the colony is plotted as a function of normalised time for the two progeny enclaves (colored red and green) and for the colony (blue) in case of cells growing at 37 *^°^*C (**A**) and 25 *^°^*C (**B**).

**Supplementary Figure 10:**
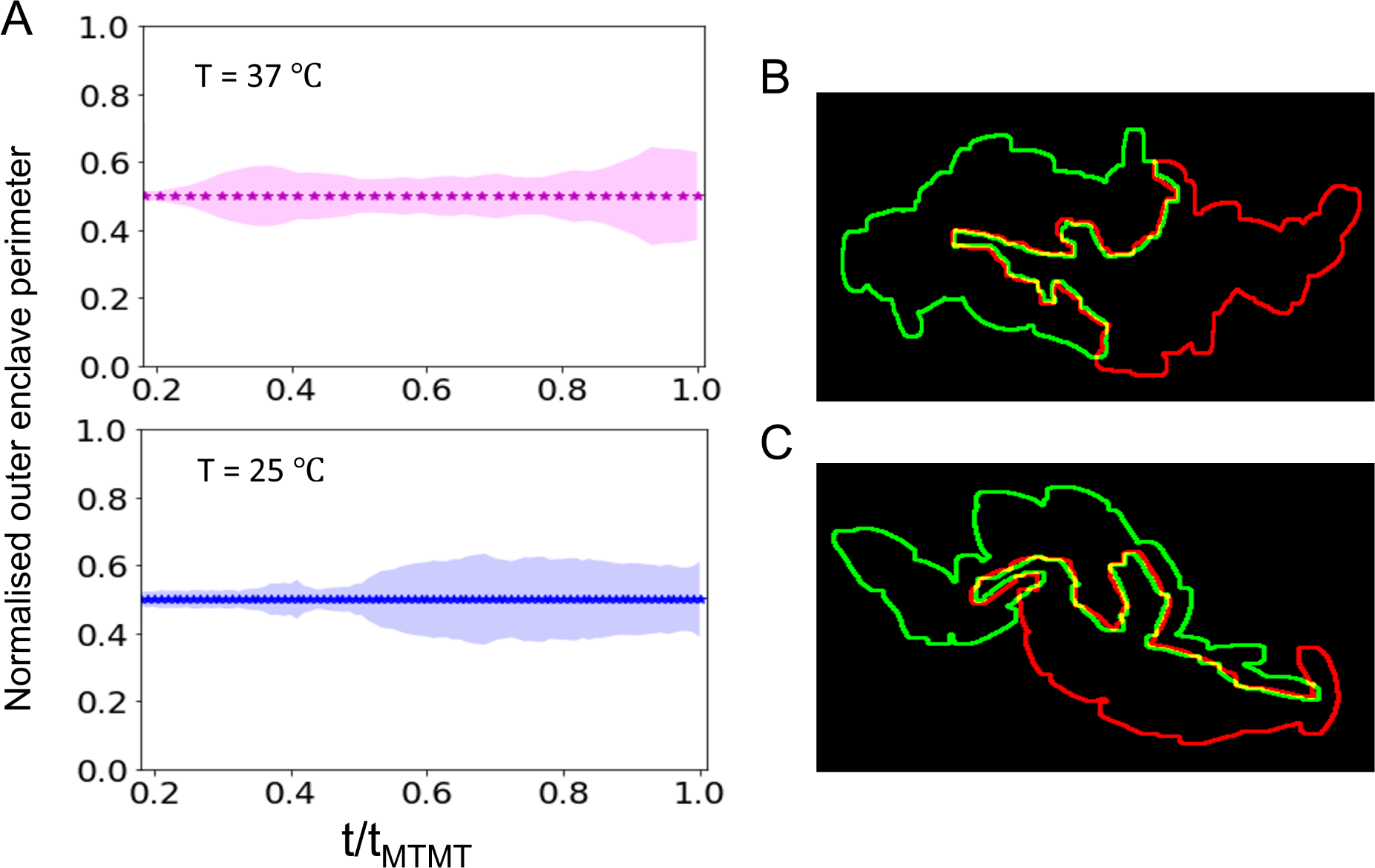
Enclaves have identical exposure to external surroundings. **A.** Enclave perimeter which is exposed to the outer surroundings (referred to as outer enclave perimeter) for each enclave is normalised by colony perimeter and plotted as a function of dimensionless time (*t/t_MT MT_*). **B,C.** show the enclave outer boundary and the enclave interface for a representative colony at 37 *^°^*C (**B**) and 25 *^°^*C (**C**).

**Supplementary Figure 11:**
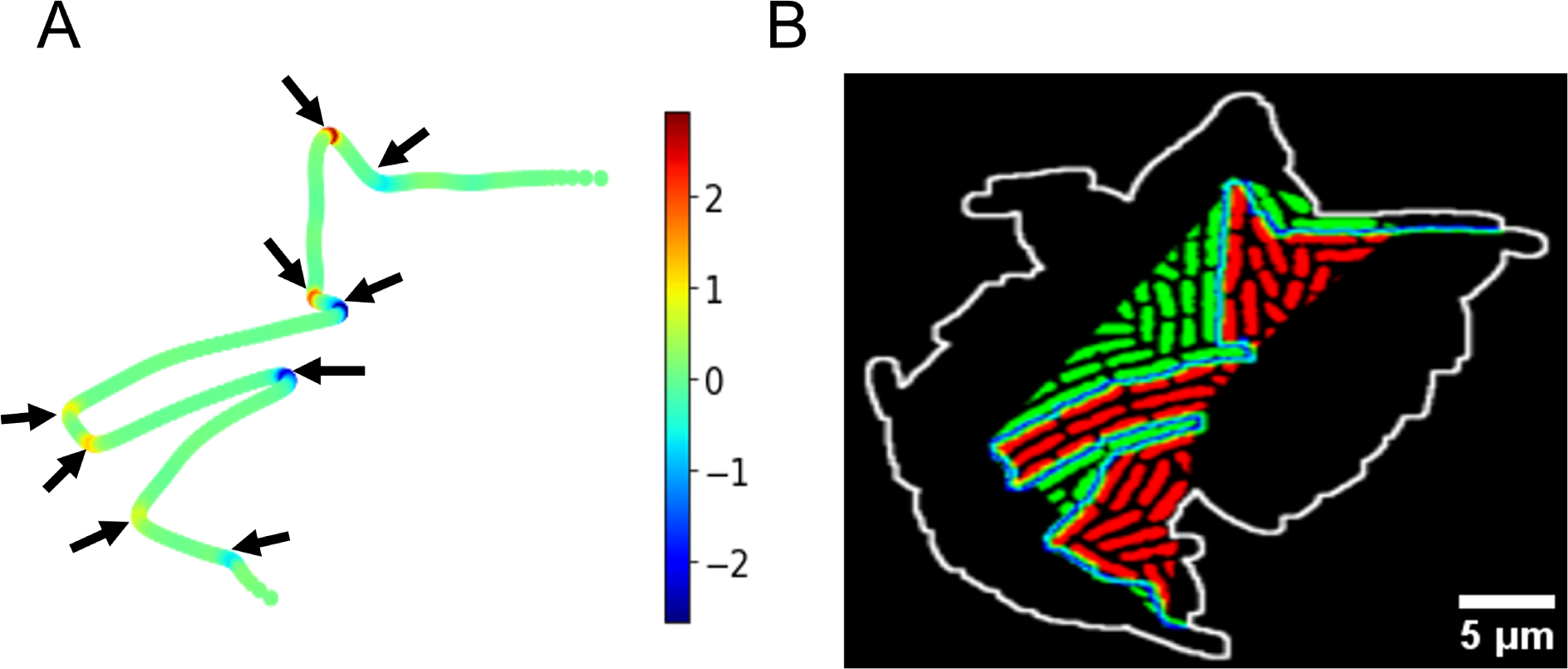
Geometry of invasion and interfacial curve. **A.** Colormap of the signed curvature of the enclave interface curve computed using kappa plugin. Black arrows mark high curvature regions on the interface. **B.** Using high curvature points which marked the beginning of invasion front, we compute the region of invasion of one enclave into other.

**Supplementary Figure 12:**
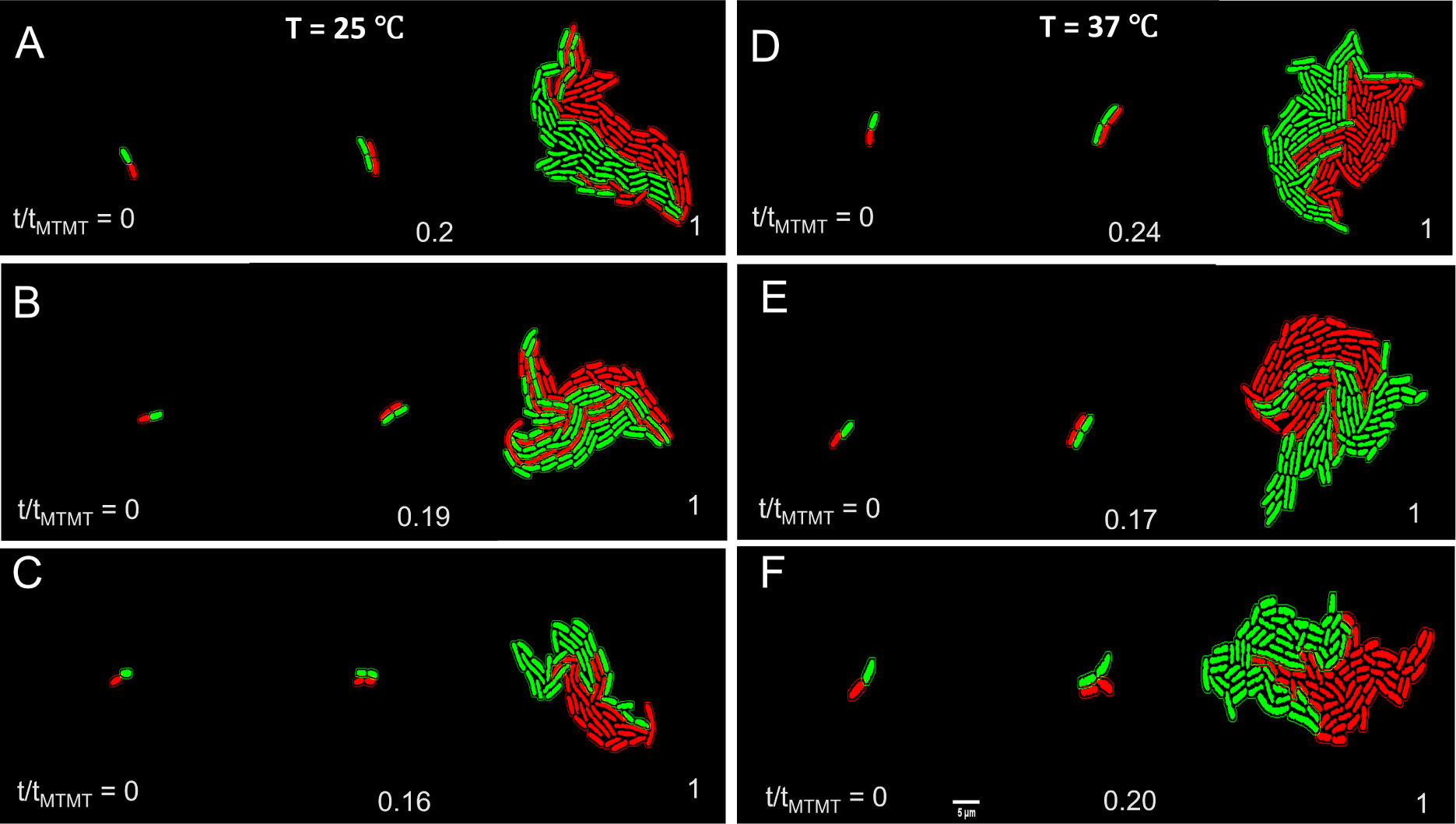
Enclave formation is independent of arrangement of cells at the early stages of colony formation. Initial arrangement (*N_r_* = 2, *N_g_* = 2) of the two descendant cells is similar for the three replicas, shown for a representative colony growing at 25 *^°^*C (**A-C**) and at 37 *^°^*C (**D-F**). The final cell arrangement geometry for both the temperatures is clearly independent of the initial arrangement, in all cases.

**Supplementary Figure 13:**
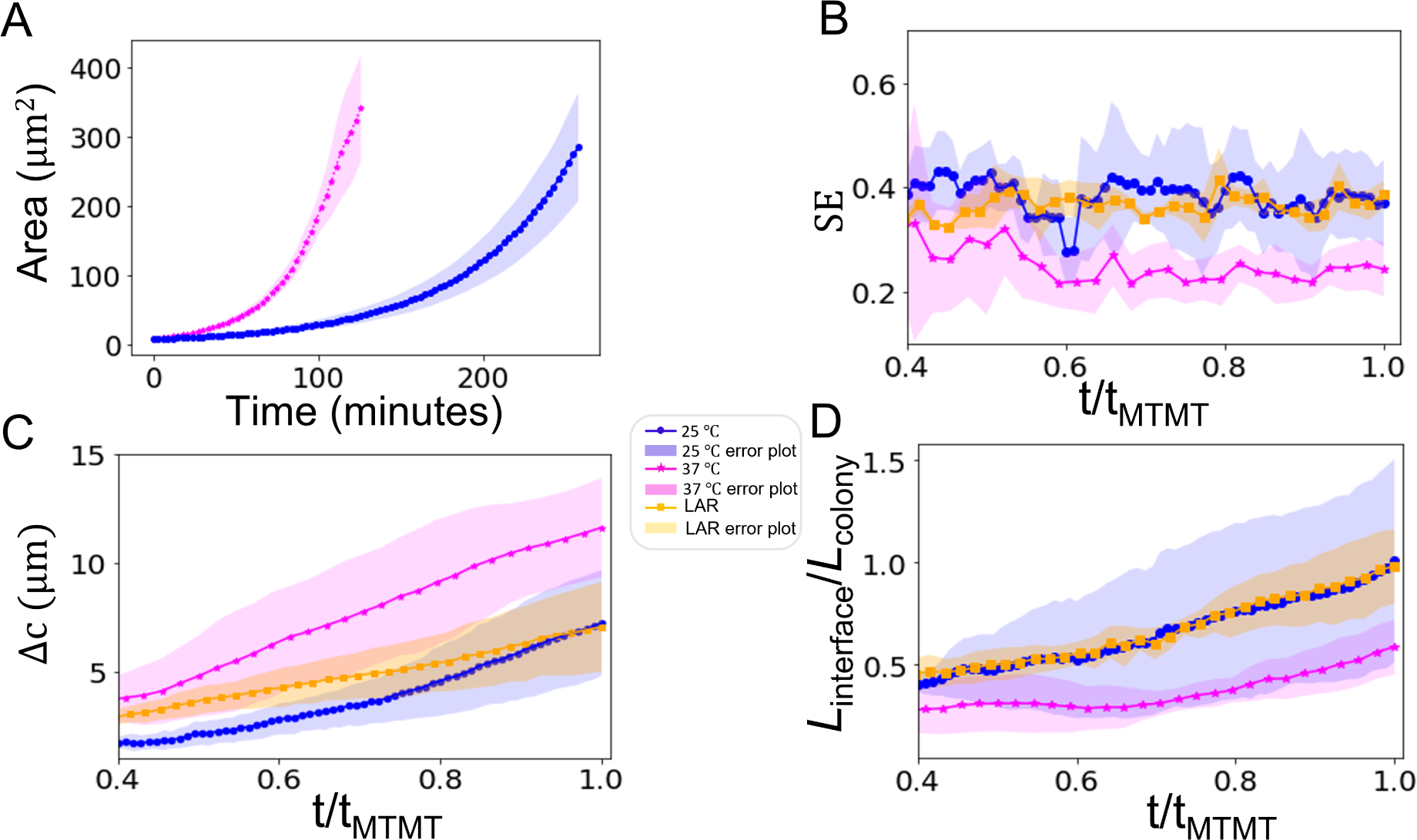
Cell arrangement patterns are consistently more disordered for slower growing cells, display a similarity with cell arrangement patterns in colonies of low aspect ratio cells. **A.** The areal growth curve of the progeny enclaves is shown for *E.coli* cells growing at 37 *^°^*C (magenta) and 25 *^°^*C (blue). **B-D.** show comparison of time series of Shannon entropy (SE), relative distance between the centroids of the two enclaves (Δ*c*) and normalised interfacial length (*L_interface_*/*L_colony_*) for *E.coli* colonies growing at 37 *^°^*C (magenta (stars)) and 25 *^°^*C (blue (circles)), and orange (squares) denote the colonies formed by LAR cells.

**Supplementary Figure 14:**
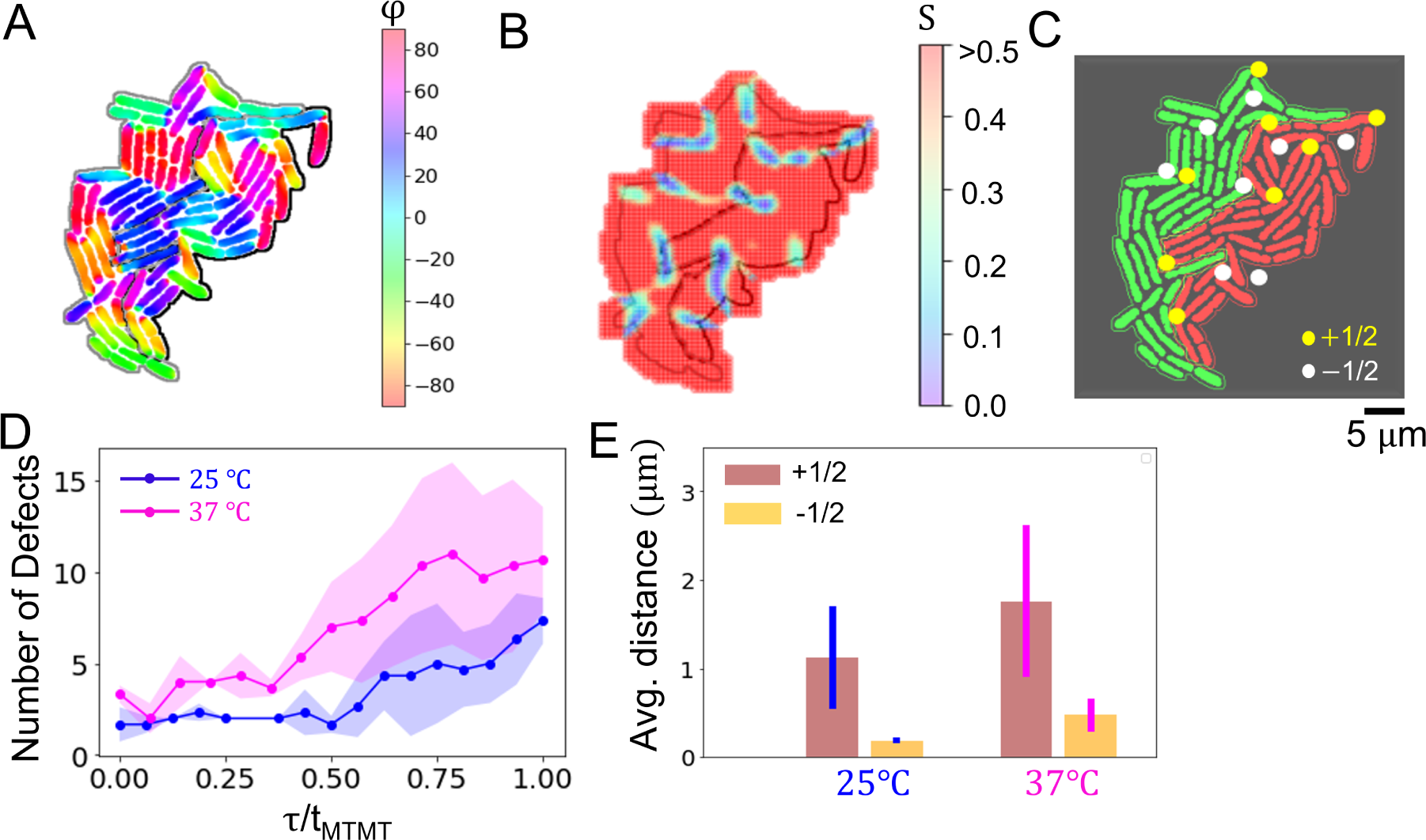
Detection and tracking of topological defects. **A.** Colormap of orientation of cells in a representative colony with the enclave interfacial curve marked in black depicting that the majority of cells in the interface are aligned tangentially, **B.** Colormap of the nematic order parameter S, **C.** Topological defects of charge +1*/*2 (yellow dots) and −1*/*2 (white dots) marked out in the colony, with progeny enclaves colored red and green, **D.** Total number of defects for colonies growing at 25 *^°^*C (blue) and 37 *^°^*C (magenta) as a function of normalised time, **E.** Average distance travelled by +1*/*2 defects and −1*/*2 defects in colonies growing at 25 *^°^*C and 37 *^°^*C.

**Supplementary Figure 15:**
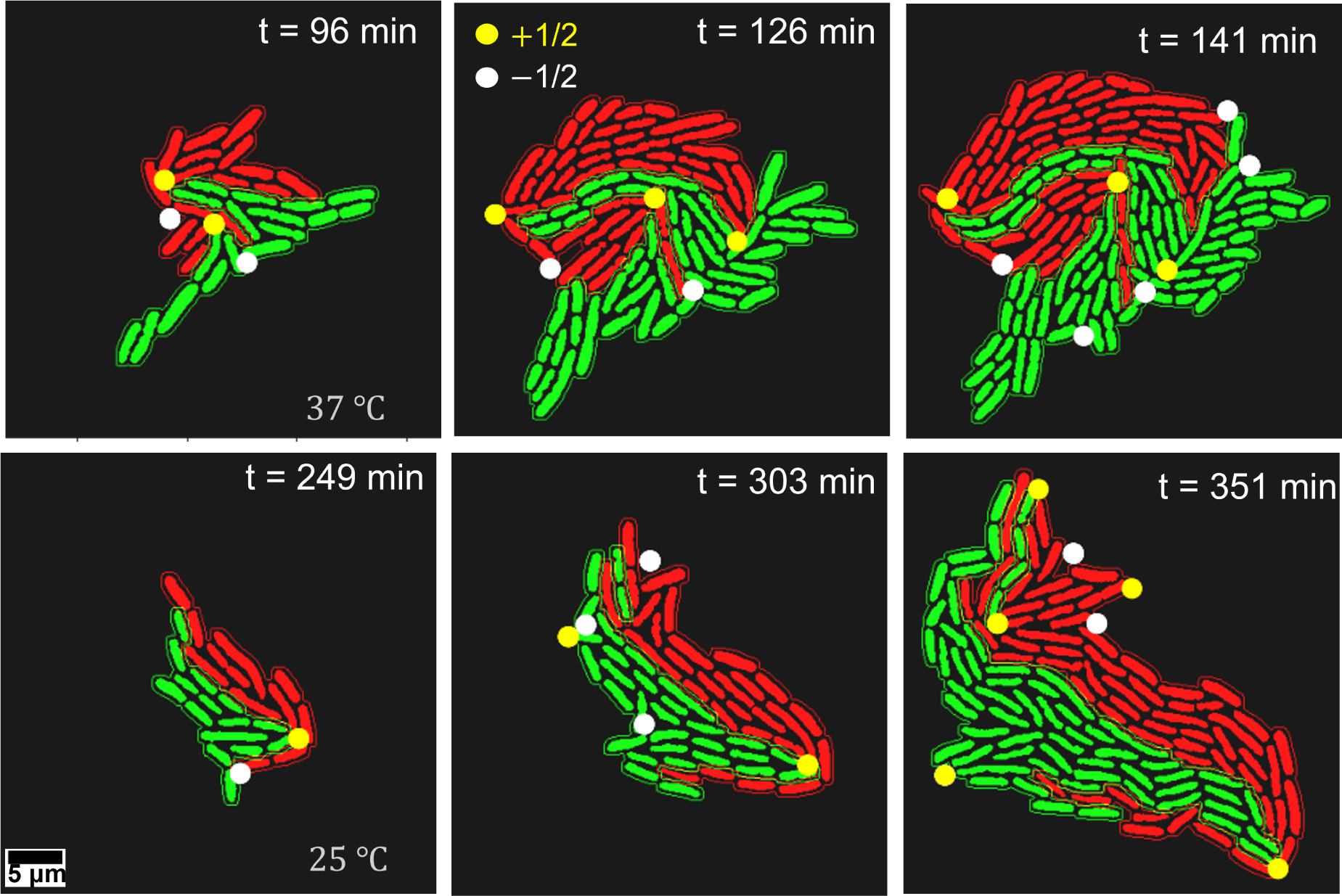
Distribution of topological defects in bacterial colonies. Topological defects in representative colonies growing at 37 *^°^*C (up) and 25 *^°^*C (down) as they evolve with time. Note the proliferation of defects along the interface boundary.

**Supplementary Figure 16:**
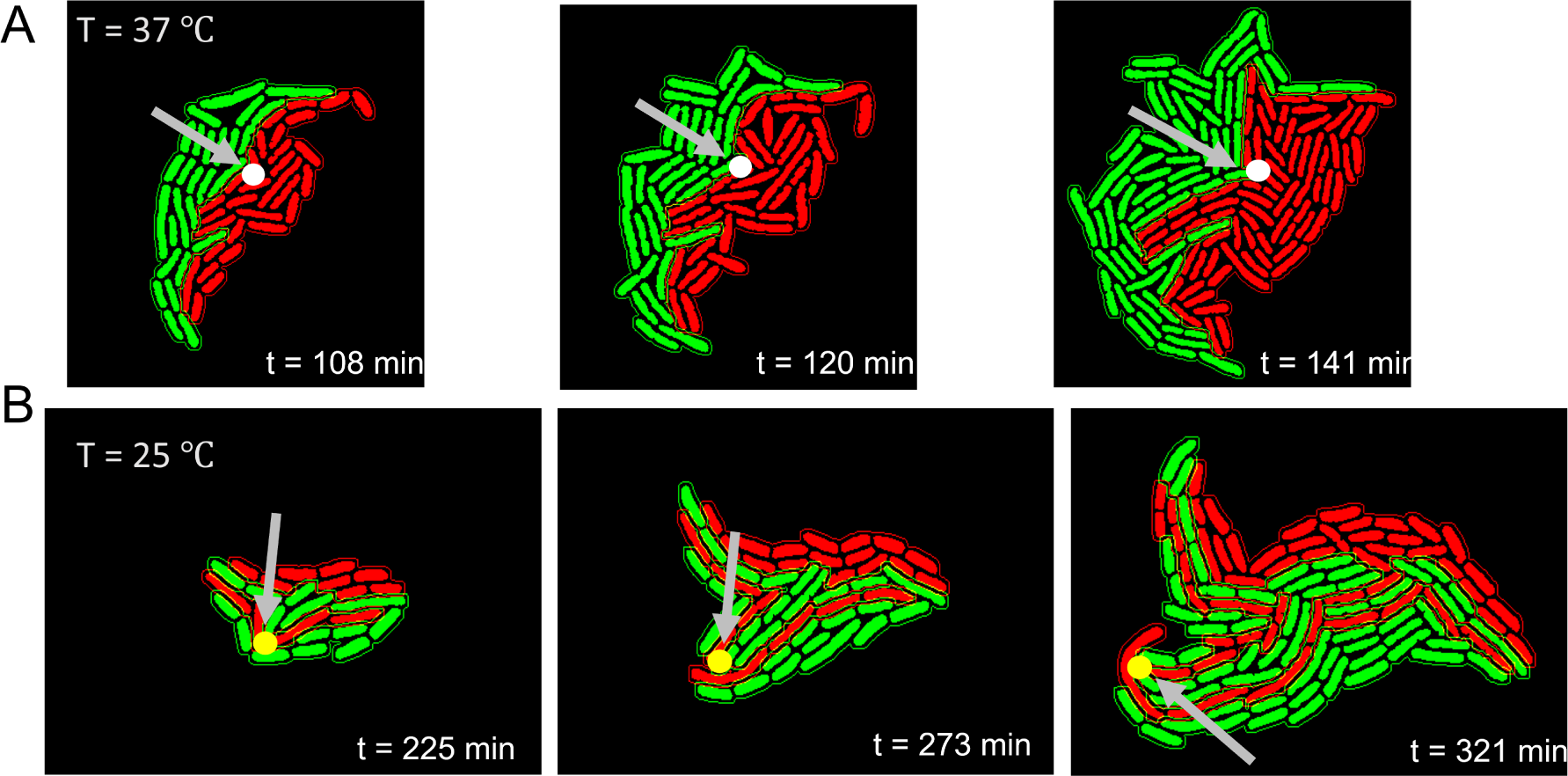
Topological defects initiate enclave invasion and persist as invasion deepens. **A.** Initiation of a wide front enclave invasion nucleated by a −1*/*2 defect, which is marked by the arrow (left), with the defect persisting as the invasion front deepens in a colony growing at 37 *^°^*C (right). **B.** Initiation of a narrow front invasion nucleated a +1*/*2 defect (left) with the defect persisting as the invasion deepens in a colony growing at 25 *^°^*C (right).

**Supplementary Figure 17:**
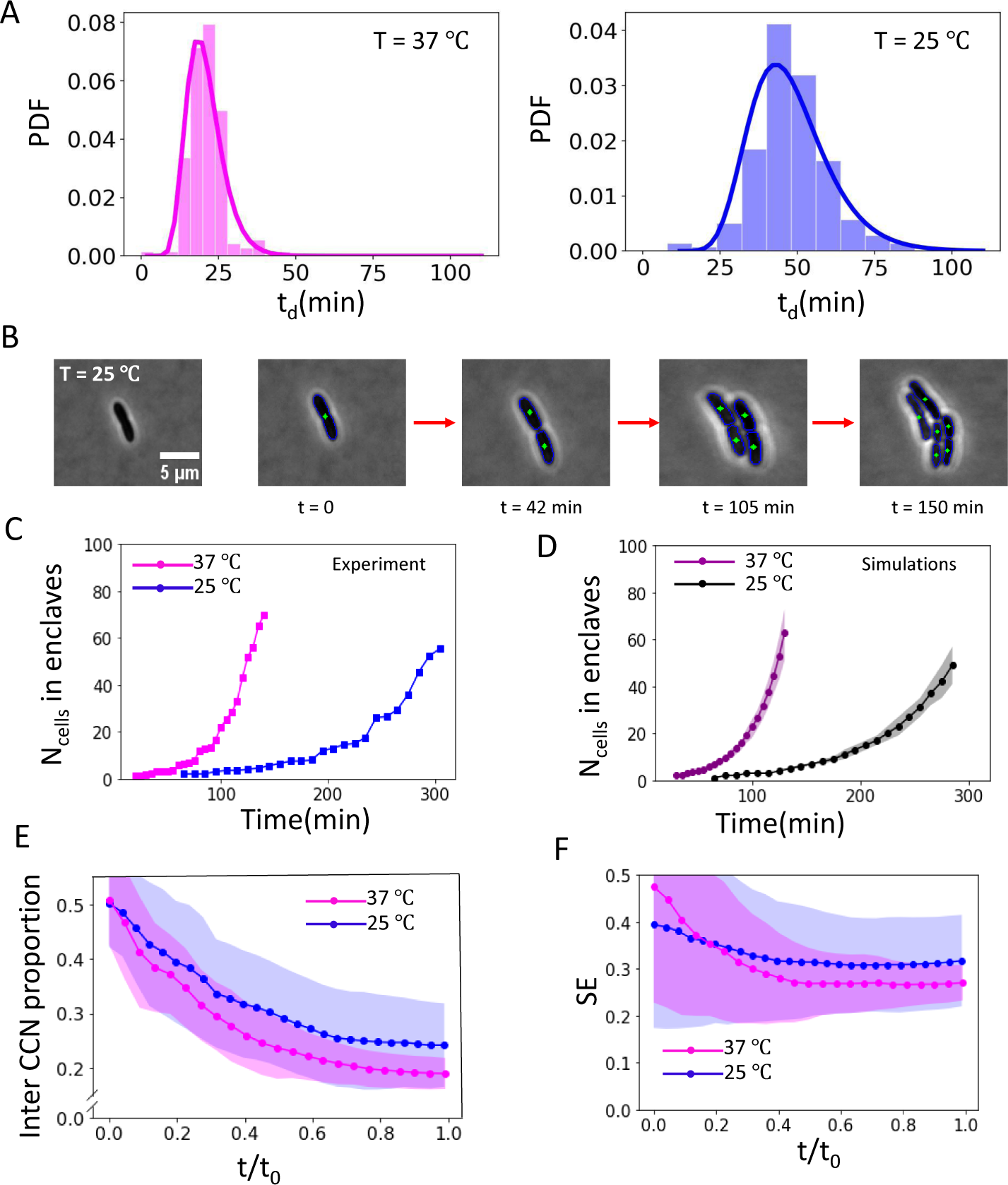
Distribution of division times of cells in colonies, cell number evolution in colonies and analysis of simulated colonies. **A.** Normalized frequency for cell division time is plotted by tracking the division times of the cells in the colony as it evolves. The mean and standard deviation of the division times calculated from all replicas is *t_d_* = 21 ± 6 minutes at 37 *^°^*C and *t_d_* = 48 ± 13 minutes at 25 *^°^*C and the data was fitted with appropriate log normal distributions. **B.** Illustrative example of division time of the founder cell and subsequent daughter cells as they grow and divide at 25 *^°^*C. **C.** Growth of number of cells in colonies growing at 25 *^°^*C and 37 *^°^*C, observed in our experiments, **D.** Growth of number of cells in simulated colonies from our lattice model simulations, See Supplementary Sec. S.9. **E.** Proportion of Inter-enclave contacts for simulated colonies as function of normalised time, **F.** Shannon entropy of arrangement patters for simulated colonies as a function of time.

**Supplementary Figure 18:**
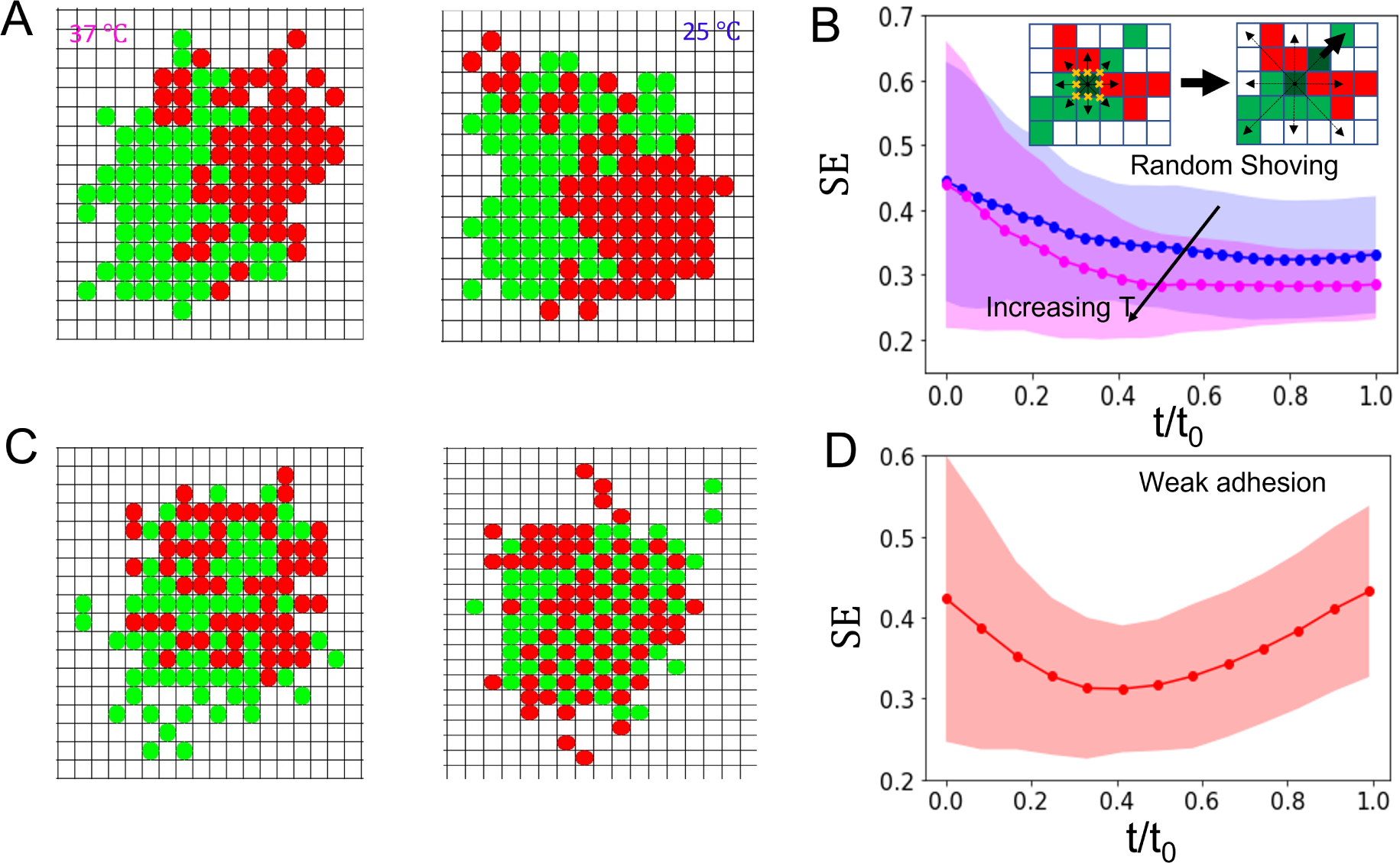
Random cell shoving model and weakened daughter cell interaction model. **A.** Colonies simulated using cell shoving where direction is chosen randomly with equal probability, **B.** The evolution of Shannon entropy of arrangement with normalised time for faster growing (magenta) and slower growing (blue) colonies (INSET)-Schematic of random cell shoving of a dividing cell whose Moore neighbourhood is filled up, **C.** Snapshots of simulated colonies where one daughter cell was allowed to be placed away from neighbourhood of the mother cell (in particular two lattice steps away), thus not requiring two daughter cells to always be neighbours, in effect weakening the adhesion of daughter cells, **D.** Evolution of Shannon entropy of arrangements with normalised time for this case.

**Supplementary Figure 19:**
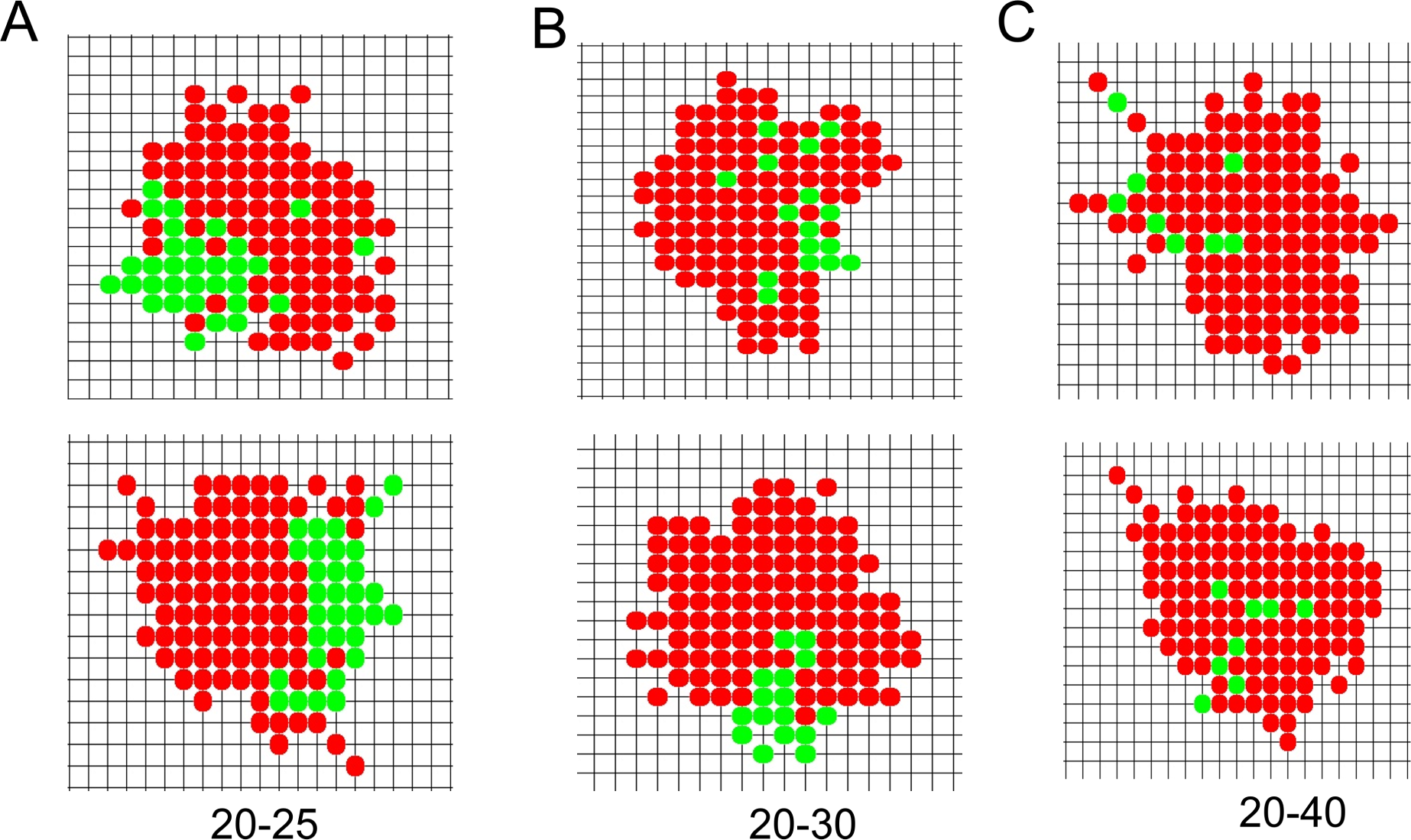
Simulations of colonies comprising cells having different division times. Lattice model simulations of colonies starting from two initial cells where one cell and its descendants divide at 20 minutes and the other cell and its descendants divide at 25 mins (**A**), 30 mins (**B**) and 40 mins (**C**). While in (**A**) formation of progeny enclaves is clear, for (**B**) in some cases enclaves are formed while in other cases much larger degree of intermixing occurs. In (**C**), enclaves are almost always absent.

## Notes

### Competing Interest Statement

The authors have declared no competing interest.

